# Challenges in Predicting Chromatin Accessibility Differences between Species

**DOI:** 10.1101/2025.11.09.687449

**Authors:** Amy Stephen, Arian Raje, Heather H. Sestili, Morgan E. Wirthlin, Alyssa J. Lawler, Ashley R. Brown, William R. Stauffer, Andreas R. Pfenning, Irene M. Kaplow

## Abstract

Enhancers are transcriptional regulatory elements that help drive phenotypic diversity, yet they often undergo rapid sequence evolution despite functional conservation, posing a challenge for predicting their function across species. Machine learning models that predict quantitative enhancer activity using DNA sequence have not previously been evaluated for their ability to predict quantitative differences across orthologous regions. Here, we trained convolutional neural networks (CNNs) on a regression task to predict chromatin accessibility, which is a proxy for enhancer activity, in the liver across five mammals, and we developed a novel framework to evaluate cross-species performance. We demonstrated that training on multiple species improves model generalization to both species used in training and held-out species. However, the models consistently achieved poor performance in predicting quantitative differences in accessibility between species at orthologous regions. Our study highlights the challenges in using regression models to predict chromatin accessibility changes between species.

## INTRODUCTION

Enhancers are transcriptional regulatory elements that loop in three dimensions to transcription start sites to activate gene expression (1). Within a given tissue, the regulatory activity of an enhancer is shaped by a combinatorial code of transcription factor (TF) binding motifs and other local sequence features (2–5). Enhancer activity can be approximated by measures of chromatin accessibility from assays like DNase-seq and ATAC-seq, as regions of the genome that are accessible tend to be easily bound by transcription factors (6, 7).

Unlike protein-coding sequences, which are often highly conserved, enhancer regions often evolve rapidly at the nucleotide level despite their essential roles in development and disease (8). Many phenotypic differences between species and even within species have been linked to changes in enhancer activity rather than changes in protein-coding genes (9–11). Understanding how enhancer function is conserved or diverges across species is critical for understanding the mechanisms underlying the gene expression differences between species.

Although many functional enhancers lack strong nucleotide-level conservation (8, 12, 13), the conservation of the presence of TF binding motifs and other local sequence features offers an alternative approach for quantifying enhancer activity conservation. Previous studies were able to train machine learning models on sequences underlying candidate enhancers to predict whether an enhancer will be active. They found that these models could make accurate predictions in species not used in training (14, 15), including at locations where the enhancer activity differed from that of the orthologous region in the training species (15, 16).

A key limitation of these models is their binary treatment of enhancer activity, which does not reflect the continuous nature of chromatin accessibility data (17, 18). Even for single-cell chromatin accessibility assays, data is typically analyzed by pooling cells that likely come from the same cell type, thus creating a quantitative enhancer activity measurement for each cell type. Many candidate enhancers are present at orthologs across multiple species but have differences in accessibility at these orthologs. Such differences in accessibility might affect gene expression and downstream phenotypes.

Previous work by Kelley (19) demonstrated that regression models can be trained to predict chromatin accessibility across species using input sequences that are over 100kb long. The study found that a model trained using data from two species performs better in both species than a model trained using data from only one. The study suggested that this performance boost is due to the increased genomic diversity of the training data rather than to the increased number of species in the training set. While this work shows the potential of using regression models for chromatin accessibility prediction trained in multiple species, the ability of the models to predict quantitative differences in chromatin accessibility between species was not evaluated. In addition, long sequences cannot be used as input in many current genome assemblies, as these assemblies often consist of scaffolds that are only a few hundred thousand base pairs long (20, 21).

Large-scale initiatives such as the Vertebrate Genomes Project (22) and the Bat1K Project (23) are in the process of building chromosome-level assemblies for hundreds of mammals, but alignments of these genomes have not yet been completed. In addition, the majority of sequenced mammals are not included and will continue to have fragmented assemblies for likely many years to come due to the difficulties of obtaining high-quality DNA from animals that live in remote locations (21, 24). Thus, training models with shorter input sequences is essential for enabling predictions in most available assemblies.

Previous work showed that classification models for enhancer activity prediction that work well on parts of the genome not used in training cannot always accurately predict gains and losses in enhancer activity in species not used in training (13, 25). In one such study, some convolutional neural network (CNN) machine learning models with impressive performance in predicting whether a region would be accessible in the brain on held-out chromosomes had poor performance in predicting the gain or loss of a brain accessible region in a new species (13). In another study, a gapped k-mer support vector machine learning model trained on human sequences from a STARR-seq assay in HepG2, a liver cell line, was unable to predict the relative STARR-seq activity of enhancers from non-human primates when normalized to their human orthologs (25).

Furthermore, recent studies have demonstrated that regression models trained to predict gene expression do not accurately predict expression differences between individuals (26, 27), further demonstrating that models trained to predict continuous genomic measurements often fail to predict how those measurements change in the presence of changes in sequence. The limitations illustrated in these previous studies demonstrate the need for systematically evaluating the ability of regression models to make accurate predictions of enhancer activity differences between species.

We addressed this need by developing a novel framework for evaluating the ability of models to predict chromatin accessibility differences between species. We then used our framework to evaluate how well multiple approaches for training machine learning models could predict differences in chromatin accessibility between species. For this evaluation, we created a new dataset for comparing chromatin accessibility between species, which involved generating new ATAC-seq data from the rhesus macaque (*Macaca mulatta*) and Norway rat (*Rattus norvegicus*) and uniformly processing it along with publicly available ATAC-seq data from the house mouse (13, 28), cow, and pig (29). We focused on the liver due to its cell type homogeneity relative to other tissues (30).

We trained our CNNs using primarily 500 base pair (bp) sequences to predict chromatin accessibility so that we could use them to make predictions at orthologous regions in fragmented assemblies. We tested multiple strategies for normalizing accessibility signals across species, multiple convolutional neural network hyper-parameters, multiple training sequence input lengths, and multiple subsets of species for model training. We found that many of the models worked well on held-out regions of species used in training and performed better than random when predicting chromatin accessibility differences between species, but none achieved strong performance on this task. Consistent with previous studies, we found that training on data from multiple species improved model performance for both species represented in the training data and for held-out species (19), but performance in predicting differences in accessibility between species was poor even for models trained using data from all five species. Our results demonstrate the importance of evaluating models for cross-species enhancer prediction for their ability to predict enhancer activity differences between species and that training a model to make such predictions accurately remains a challenge.

## MATERIALS AND METHODS

### Animal Use and Sample Collection

We obtained tissue samples from corresponding tissues from two rhesus macaques and three Norway rats. Specifically, we obtained liver and multiple brain regions. We chose to focus our analysis on data from the liver samples for this study because recent work has suggested that brain regions may have different cell type proportions in different species (31–33).

We performed all animal procedures in accordance with the National Institutes of Health Guide for the Care and Use of Laboratory Animals and approved by the Institutional Animal Care and Use Committees (IACUC) of Carnegie Mellon University (Protocol ID 201600003) and the University of Pittsburgh (Protocol ID 19024431). We single- or pair-housed rhesus macaques at the University of Pittsburgh with a 12h-12h light-dark cycle. The macaques in this study were a 12-year-old female (8.1 kg) and a 4-year-old male (6.0 kg). Before surgery, we sedated the macaques with ketamine (15 mg/kg IM) and then ventilated and further anesthetized them with isoflurane. We next transported the animals to a surgery suite and placed them in a stereotaxic frame (Kopf Instruments). We opened the body cavity and excised a sample from the right lobe of the liver. Separately, we removed the calvarium and then perfused the circulatory systems with 3-4 liters of ice cold, oxygenated macaque artificial cerebrospinal fluid (124 mM NaCl, 5 mM KCl, 2 mM MgSO4, 2 mM CaCl2, 3 mM NaH2PO4, 23 mM NaHCO3, 10 mM glucose). We then opened the dura and removed the brain. We sliced the brains into coronal slabs using a brain dissection knife. We excised the following brain regions under a dissection microscope: secondary auditory cortex (2A), premotor area 6V (6V), supplementary motor area (SMA), hand and forearm motor cortex (hM1), hand and forearm S1 (hS1), orofacial motor cortex (ofM1), orofacial S1 (ofS1), cerebellum, caudate, nucleus accumbens (NAcc), and putamen. We also collected liver, primary motor cortex (M1), and caudoputamen of the striatum (STR) tissues from three rats (1 male Sprague-Dawley, housed in the University of Pittsburgh; 2 Brown Norway, 1 male and 1 female, housed at Carnegie Mellon University) (**Supplementary Fig. S1**).

We euthanized rats by isoflurane overdose followed by decapitation. We collected livers immediately and kept them cold on ice in lysis buffer (6). We sliced brains into 300 μm sections in a vibrating microtome (Leica VT 1200) in ice-cold, oxygenated rodent artificial cerebrospinal fluid [119 mM NaCl, 2.5 mM KCl, 1 mM NaH2PO4 (monobasic), 26.2 mM NaHCO3, 11 mM glucose]. We sampled the brain regions from these coronal sections under a dissection microscope and transferred them to a chilled lysis buffer (34).

### ATAC-seq

We processed tissue samples as described previously (6, 34) with the following minor differences in procedure and reagents: We isolated nuclei from dissected tissues using 30 strokes of homogenization with the loose pestle (0.005 in. clearance) in 5mL of cold lysis buffer placed in a 15 mL glass Dounce homogenizer (Pyrex #7722-15). We filtered the nuclei suspensions through a 70 µm cell strainer, pelleted them by centrifugation at 2, 000 x g for 10 minutes, resuspended them in water, and filtered them a final time through a 40 µm cell strainer. We stained sample aliquots with DAPI (Invitrogen #D1206) and quantified nuclei concentrations using a manual hemocytometer under a fluorescent microscope. We input approximately 50, 000 nuclei into a 50 µL ATAC-seq tagmentation reaction as described previously (6, 34). We amplified the resulting libraries to 1/3 qPCR saturation and estimated fragment length distributions using the Agilent TapeStation System. We shallowly sequenced barcoded ATAC-seq libraries at 600, 000-5 million reads per sample on an Illumina MiSeq and processed individual samples through the ENCODE pipeline (35), including running Preseq to estimate library complexity (36). We used the quality control measures from the pipeline to filter out the lowest-quality samples and re-pool an approximately balanced library for paired-end deep sequencing. We did the deep sequencing on an Illumina NovaSeq 6000 System, where we used the University of Pittsburgh sequencing core for the first macaque and the first rat and GENEWIZ’s services for the other animals.

### Constructing positive and negative sets

We used the bulk ATAC-seq data that we collected from the liver of the rhesus macaque (*Macaca mulatta*) and Norway rat (*Rattus norvegicus*) as well as publicly available bulk liver ATAC-seq data from house mouse (*Mus musculus*) (15, 28), cow (*Bos taurus*), and pig (*Sus scrofa*) (29). A summary can be found in **Supplementary Table S1**. We processed the datasets as we described in (13) (**Supplementary Table S2**). This involved running the ENCODE ATAC-seq pipeline (35), which uses bowtie2 for mapping reads (37), picard (38) and SAMtools (39) for filtering reads, MACS2 for calling peaks (40), and the Irreproducible Discovery Rate (IDR) for finding reproducible peaks (41). Since, for each animal, our samples had substantial differences in sequencing depth, we used the IDR reproducible peaks across pooled pseudo-replicates.

We used the MACS2 peak caller “fold_enrichment” column value as a quantification of chromatin accessibility. Higher MACS2 signal values reflect a higher proportion of cells in which a given regulatory region is accessible. Specifically, the signal in MACS2 is the ratio of read count to the local estimation of background noise within each peak (40). Although MACS2 does not claim to explicitly normalize for region length in its read count calculations, ATAC-seq peaks are generally of similar lengths on the order of hundreds of base pairs (42, 43), particularly following our filtering step in which we excluded all regions greater than 1 kb (13). As a result, signal strength estimates from MACS2 should be approximately comparable across peaks and species in our ATAC-seq datasets.

We found that many of the new datasets we generated were high-quality; full data quality reports can be found in the **Data Availability** associated with this manuscript. We used all of the liver samples except for macaque biological replicate 2, which had poor periodicity, and rat biological replicate 1, which had less than 90% of reads properly paired in both technical replicates. We should also note that the publicly available cow and pig data had poor periodicity and that the pig data had less than 90% of reads properly paired in both biological replicates. We included them because we previously found that classification models trained using other datasets made accurate predictions for these (13). In addition, the pig genome assembly we used (susScr3) was of lower quality than the rat genome assembly we used (rn6), so the lower fraction of properly paired reads from the pig is less likely to be indicative of poor data quality.

To identify candidate enhancers in each species and use them to create our positive and negative sets, we used the process described in our previous work (13). Since we intersected reproducible peaks from two datasets to obtain the mouse liver candidate enhancers, we used the signal value from the peaks generated in our previous work because they were collected from fresh instead of flash-frozen tissue (13). For model training, we set the signal values for the negative set examples to zero. We evaluated results separately on positive and negative sets to determine the extent to which our performance was caused by the model accurately quantifying chromatin accessibility and not just differences between positives and negatives. We used two examples for each sequence – one with the sequence and another with its reverse complement. We split our training, validation, and test sets as described in our previous work (13, 15, 44). For training models using only mouse sequences, we used only the mouse training and validation set; for training models using sequences from mouse, rat, and macaque, we used the training and validation sets from all three of these species; for training models using sequences from mouse, rat, macaque, cow, and pig, we used the training and validation sets from all five of these species.

### Quantile normalization

We initially trained our models on log-transformed signal values but were concerned that MACS2’s normalization strategy may be insufficient for cross-species comparisons, as signal distributions varied slightly between species (**Supplementary Fig. S1A**). To address this, we trained models on quantile normalized data. Quantile normalization normalizes signal values across datasets by ranking samples and then giving all samples at each rank the average signal value across all datasets (45). Since quantile normalization requires equal sample sizes, we selected the peaks with the strongest signal, hypothesizing they would be most informative for model training. We first log-transformed chromatin accessibility signals, next selected the strongest 20, 615 peaks from each species (the number of peaks in the species with the smallest number of reproducible peaks that were candidate enhancers after filtering), and then ranked peaks within each species (rank 1 for the highest signal, rank 20, 615 for the lowest). For each rank *i*, we averaged the log-transformed signal strength across all five species. We then assigned the average as the normalized signal for rank *i* in each species (**Supplementary Fig. S1B**).

### Extended quantile normalization

One limitation of quantile normalization is that it requires equal sample sizes, and we had different numbers of reproducible peaks for each species (**Supplementary Table S2**). To address this, we developed a novel method called extended quantile normalization (EQN). This approach applies quantile normalization to a random subset of peaks equal to the number of peaks from the smallest sample. It then assigns normalized values to the remaining peaks by matching each to its nearest neighbor (in log-transformed signal space if using log-transformed signal) within the quantile-normalized subset. This strategy allows us to use the full dataset while obtaining comparable signal distributions across species, with values reflecting relative signal strengths (**Supplementary Fig. S1C**).

### Cross-species evaluation set creation

We generated validation and test sets that allowed us to directly evaluate our model’s ability to predict differences in chromatin accessibility between species (**Fig. 1**). We did this using bedtools v2.30.0 (46). To explain validation and test set generation, we will describe the process for macaque as an example. Note that we did this for each non-mouse species.

**Figure 1.**
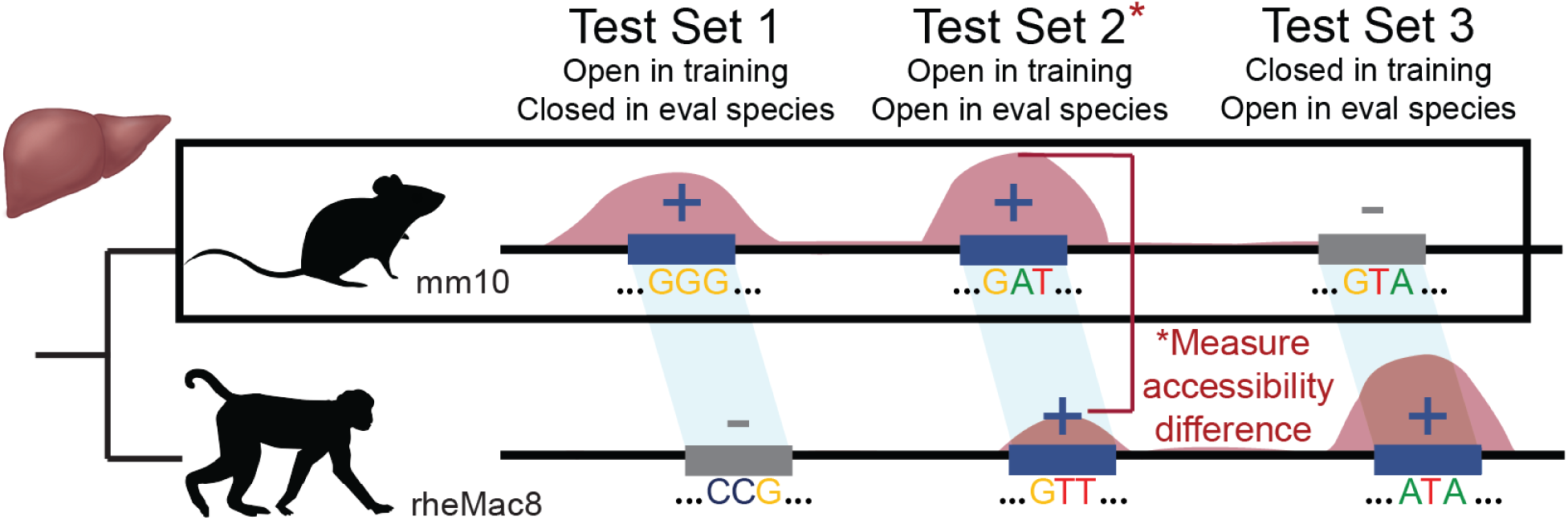
The evaluation framework is sectioned into three distinct test sets. Test set 1: regions in the evaluation species that are not accessible whose orthologs are accessible in the training species. Test set 2: regions in the evaluation species that are accessible in both the evaluation and training species. Within test set 2, we also evaluate the accessibility differences between orthologous regions as denoted by the red star. Test 3: regions in the evaluation species that are accessible but not accessible in the training species.

To generate Validation/Test Set 1, we used bedtools subtract with the -A flag to remove all macaque peaks (both reproducible and non-reproducible; we used peaks called in the pool of replicates for species with multiple replicates) from the set of mouse reproducible peaks mapped to the macaque genome. This produced a set of macaque non-accessible regions whose orthologs are candidate enhancers in the mouse. To generate Validation/Test Set 2, we used bedtools intersect with the -wa -wb flags to identify unique macaque test set peaks mapped to mouse whose orthologs overlapped with mouse reproducible candidate enhancer peaks. This yielded the set of mouse orthologs of macaque test set peaks whose orthologs are candidate enhancers in the mouse. We then filtered the macaque peaks using this list to produce Validation/Test Set 2. To generate Validation/Test Set 3, we used bedtools subtract with the -A flag to remove all mouse candidate enhancers (both reproducible and non-reproducible) from the set of macaque test set peaks mapped to the mouse. This resulted in mouse orthologs of macaque test set peaks whose orthologs are not candidate enhancers in the mouse. We filtered the macaque peaks using this list to produce Validation/Test Set 3.

Within Validation/Test Set 2, we further evaluated differences in accessibility between orthologous regions, which required establishing one-to-one correspondence between macaque and mouse peaks. To obtain these corresponding orthologs, we used the output of the bedtools intersect -wa -wb step in the Validation/Test Set 2 creation, which was the set of macaque validation/test candidate enhancers whose mapped orthologs overlapped with reproducible mouse candidate enhancers. We created a list of one-to-one matched macaque and mouse candidate enhancers using the following rules: If a mapped macaque candidate enhancer uniquely overlapped with a single mouse candidate enhancer, we considered them a match. If one macaque candidate enhancer’s mouse ortholog overlapped multiple mouse candidate enhancers, we considered the match to be with the mouse peak with the largest overlap. If multiple macaque candidate enhancers’ mouse orthologs overlapped a single mouse candidate enhancer, we considered the match to be the macaque candidate enhancer whose mouse ortholog had the largest overlap. In cases of identical overlap sizes, we considered the match to be the candidate enhancer orthologs with the highest signal value.

To test how our models generalized to unseen clades of the phylogenetic tree, we also used the test set of laurasiatheria-specific enhancers and non-enhancers that we described in our previous work (13, 44). In brief, the positives were cow enhancers whose pig orthologs were accessible in the pooled reads across pig replicates but whose mouse, macaque, and rat orthologs were not accessible in the pooled reads across mouse, macaque, and rat, respectively. The negatives were mouse enhancers whose macaque and rat orthologs were accessible in the pooled reads across macaque and rat replicates, respectively, and whose cow and pig orthologs were not accessible in the pooled reads across cow and pig replicates, respectively.

### Training regression models

Although there is publicly available chromatin accessibility data from multiple tissues, we used only data from the liver for multiple reasons. First, the majority of the liver consists of hepatocytes (30), while other tissues are much more heterogeneous with cell type proportions varying across species (47). In addition, while related studies used multi-task models trained on large numbers of cell lines and tissues (16, 19), multi-task models have been shown to have poor performance for predicting or cell type-specific accessibility relative to single-task models, and most chromatin accessibility is tissue- or cell type-specific (48, 49).

We used convolutional neural networks (50) as our machine learning models because they have excelled at similar tasks (13, 19, 51). They can learn combinatorial sequence patterns and pertinent features, including features that were not previously known to be relevant to the biological task. Not all sequence patterns and pattern combinations involved in liver chromatin accessibility are known. Our inputs were one-hot encoded 500 bp sequences, and our outputs were continuous numerical values representing predicted log-transformed signal values, the quantile-normalized log-transformed signal values, or the extended quantile normalized log-transformed signal values. Our loss function was the mean squared error. We trained the model using the following codebase https://github.com/pfenninglab/cnn_pipeline.

We began training each model with the model architecture for the best classification model trained using mouse only data published in our previous work (This model predicted motor cortex enhancer activity.) (44). For each model type, we ran an extensive 100-model hyperparameter sweep varying the following hyperparameters: number of filters per convolutional layer, number of filters per fully connected layer, convolutional layer dropout, fully connected layer dropout, convolutional layer L2 regularization, fully connected layer L2 regularization, initial learning rate, number of convolutional layers, and number of fully connected layers. Each convolutional layer had filters with width 7 and stride 1. Each layer had a ReLU activation function except for the last layer, which had a linear activation function. Between the convolutional and fully connected layers was a max-pooling layer with size 26 and stride 26. We initialized filter weights with He-normalization (52). We used the Adam optimization method (53) with batch size 100. We trained each model until there were 4 consecutive epochs with no improvement on the validation set. The hyperparameters of the best models for each model type are listed in **Supplementary Tables S3** and **S4**. We selected hyperparameters based on their performance on validation set chromosomes in the training species, performance on Validation Sets 1-3, and performance in their ability to predict the differences in chromatin accessibility between species for Validation Set 2. After selecting the hyperparameters for each type of model, we evaluated the performance on all of the test sets.

### Motif enrichment analysis

We used motif enrichment to investigate the regulatory code between regions with different levels of chromatin accessibility in mouse. We divided mouse accessible regions based on their chromatin accessibility levels. We defined strong candidate enhancers as those with log-transformed signal levels above 2.5 (20.8% of peaks), moderate candidate enhancers as those between 1.4 and 2 (35.2% of peaks), and weak candidate enhancers as those below 1.4 (15.1% of peaks). We removed candidate enhancers with signal levels at the border of moderate and strong. We then ran differential motif enrichment between the strong and weak candidate enhancers, between the strong and moderate candidate enhancers as well as motif enrichment on all 3 groups. To do this, we used bedtools getfasta (46) to retrieve the fasta sequences for summits of the strong, moderate, and weak candidate enhancers +/- 250 base pairs (bp). We used MEME-ChIP version 4.12.0 (54) with the JASPAR 2018 database (55) to identify enriched motifs.

## RESULTS

### Datasets and model training

We generated ATAC-seq data for the liver and multiple brain regions from rhesus macaque and Norway rat. The majority of samples were high-quality, but several had poor periodicity; full quality control evaluations can be found in **Data Availability.** We excluded the brain samples from our investigation, instead focusing on liver tissue due to its much more homogeneous cell type composition (30). To investigate the ability of regression models to predict chromatin accessibility, we supplement this data with publicly available bulk liver ATAC-seq data from house mouse, cow, and pig (**Materials and methods**). We trained models on distal accessible regions (**Materials and methods**), where, for most models, we used 500 bp centered on accessible chromatin peak summits. We used 500 bp regions because they are short enough to often be mappable to species with fragmented assemblies (24), have previously been shown to be long enough to capture enough local sequence information to predict whether a region is accessible (13, 16), and are close to the length of many ATAC-seq peaks. We used the MACS2 peak caller “fold_enrichment” column to quantify chromatin accessibility, where the log of these quantifications were the target prediction values for most of the models.

### New model evaluation criteria

We developed a set of model evaluation criteria specifically for evaluating the ability to predict differences in chromatin accessibility between species. Specifically, we created multiple criteria for evaluating how well a model trained on a strict subset of species performs on datasets from other “evaluation” species, as this is necessary for determining if a model can make reliable predictions in a genome for which chromatin accessibility measurements are not available. First, as was done in previous work (19), we evaluated our models on held-out validation/test set sequences from the training species. In addition to this validation/test set (**Materials and methods**), we created the following three validation/test sets in species not used in training: regions not accessible in the evaluation species whose orthologs are accessible in the training species (Validation/Test Set 1), regions accessible in both the evaluation and training species (Validation/Test Set 2), and regions accessible in the evaluation species but not accessible in the training species (Validation/Test Set 3). For Validation/Test Set 2, we also evaluated models’ abilities to predict accessibility differences between orthologous regions.

### Strong performance on held-out chromosomes within a species does not guarantee generalization to species not used in training

We did an extensive hyperparameter search (**Materials and methods**) and obtained multiple models with strong performance on mouse chromosomes not used in training or hyperparameter tuning (**Supplementary Table S3**). Our best model trained on only mouse data had comparable performance on the training, validation, and test sequences (**Supplementary Table S5**), suggesting that the model was not overfit to the training data. We then evaluated the performance of the best model trained on only mouse data in rat, macaque, cow, and pig using our new evaluation criteria. The model successfully predicts negative instances, as their prediction distribution is skewed left towards the target value of 0 (**Supplementary Table S6** and **Fig. 2**). The moderate overlap between the distributions of predictions for positives and negatives distributions indicates that the model would function sub-optimally as a binary classifier. However, the clear separation between the prediction means and the significantly different distributions suggests that the model has learned meaningful features that distinguish the two classes to some degree, even if the boundary is not distinct.

**Figure 2.**
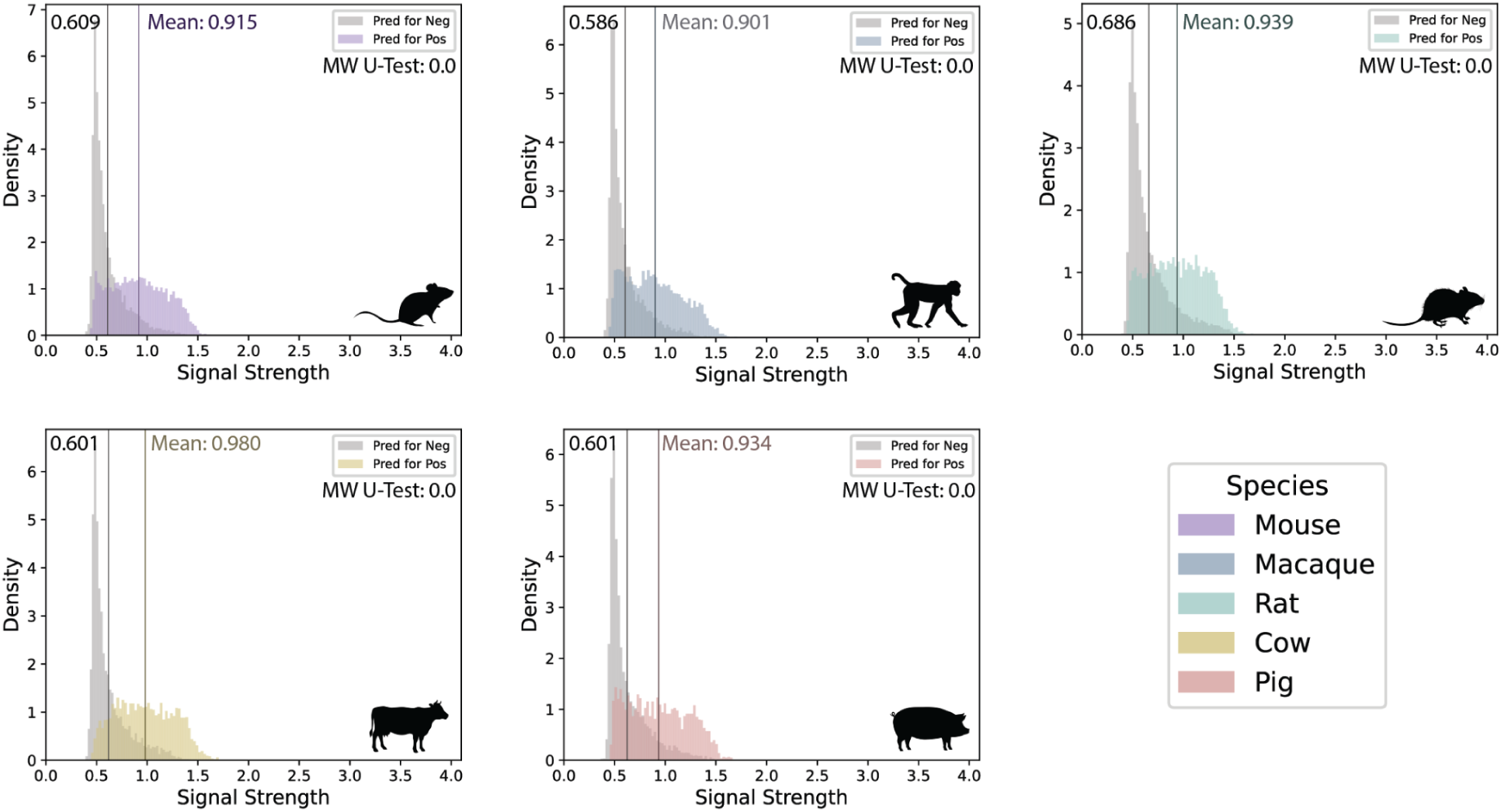
The distributions of the model’s predicted signal values are shown for the negative (gray) and positive (colored) examples. A Mann-Whitney *U* test comparing the positive and negative distributions confirmed a statistically significant difference in all cases (*p* value < 10^-30^)

Evaluation set performances for the non-mouse species were worse than for held-out mouse chromosomes. Test Set 3 performance tended to be worse than the overall test set performance for rat, cow, and pig, suggesting that the model was less reliable for predicting species-specific quantitative accessibility in species distantly related to the training species (**Supplementary Table S5**). This demonstrates the importance of evaluating chromatin accessibility prediction regression models in species not used in training.

Within Test Set 2, performance was especially poor for predicting differences in accessibility between mouse and species not used in training (**Fig. 3A**). In fact, almost half of the differences were predicted in the incorrect direction (**Supplementary Table S7**). For example, the Pearson correlation coefficient (*r*) between the predicted and real accessibility differences for mouse and cow was only 0.20, and the Spearman correlation coefficient (*ρ*) was only 0.23. This demonstrates that models with strong performance on predicting chromatin accessibility in held-out chromosomes cannot be assumed to make accurate predictions of chromatin accessibility differences between species.

**Figure 3.**
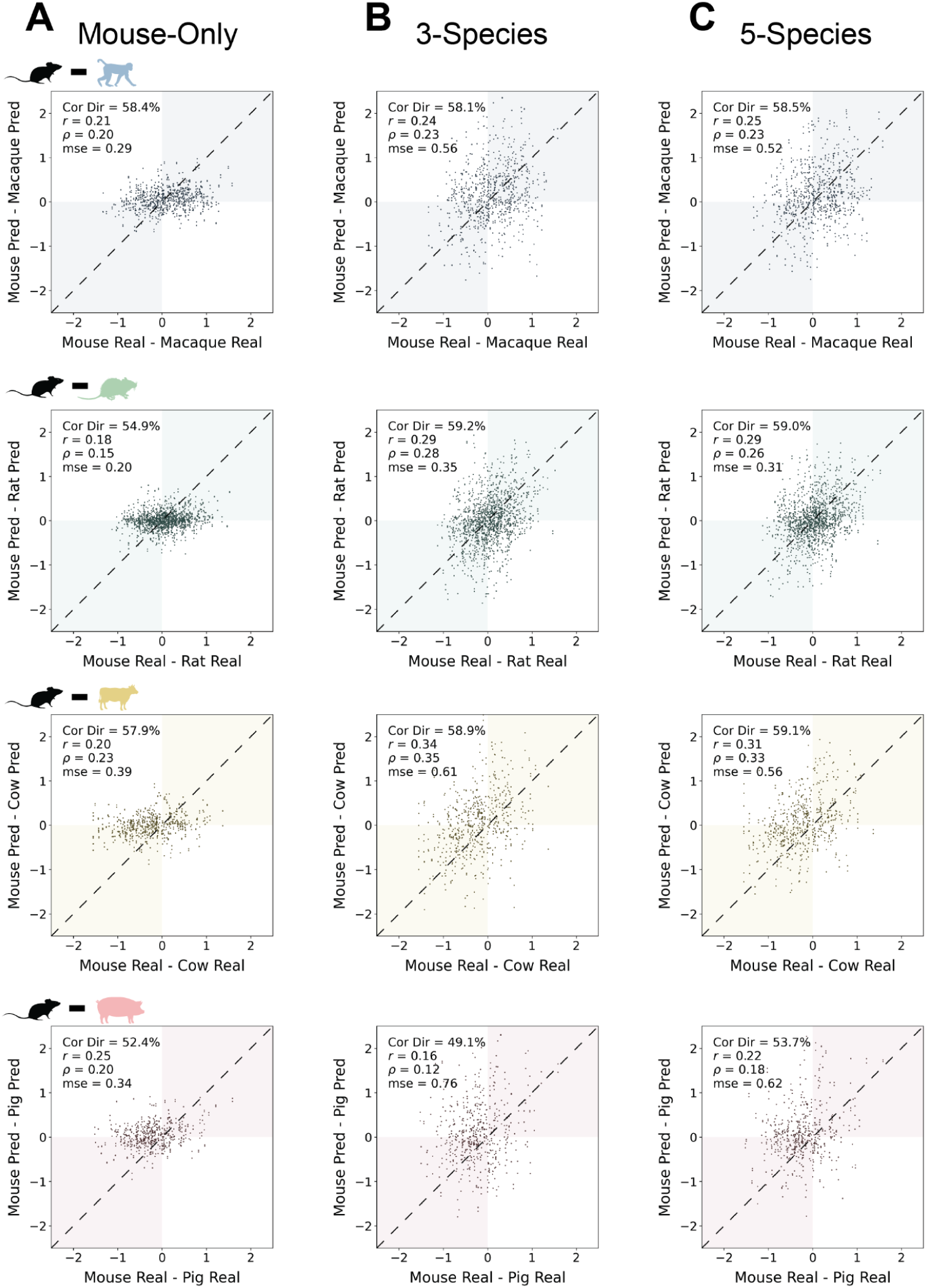
Scatter plot comparing the chromatin accessibility difference between mouse and non-mouse species of the real versus predicted signal values. Each point represents a test set 2 example (A) Chromatin accessibility difference evaluation for mouse-only trained model. (B) for 3-species model (trained on mouse, macaque, and rat) (C) for 5-species model (trained on mouse, macaque, rat, cow, and pig). Metrics reported are the percent of peaks predicted in the correction direction (Cor Dir), Pearson correlation coefficient (*r*), Spearman correlation coefficient (*ρ*), and mean squared error (mse).

We previously found that classification models with comparable performance on species used in training vary greatly in their ability to predict binary differences in accessibility between species (13). Since we had multiple sets of hyperparameters with strong performance on held-out mouse chromosomes, we evaluated models trained with different hyperparameters. Unfortunately, when evaluating these models with our new criteria, the performance was similar to that of the best mouse-only model.

### Normalizing signal across species does not improve performance

Obtaining datasets of the same read depth across species was not feasible for us (**Supplementary Table S2**) and is not feasible for most researchers due to the challenges of obtaining tissue samples from non-model organisms. Since MACS2 does not claim to explicitly normalize for total read depth, we thought that normalizing the signal across species might improve generalization to new species. Although the log-transformed signal distributions (**Supplementary Fig. 2A**) are similar across species, there may be differences preventing a model trained in one species from making accurate predictions in another. Thus, we quantile-normalized the strongest log-transformed signal values and then trained and evaluated models for predicting these quantile-normalized signals.

We found that training models on mouse quantile-normalized data generally hurt performance (**Supplementary Table S8**), with model predictions exhibiting lower correlations across all species. Notably, in the pig test set 2, *ρ* decreased by 0.08 from 0.34 to 0.26. The one exception was that the model trained to predict quantile-normalized signals more accurately predicted that negatives had signals closer to 0. Despite identical dataset sizes, the decrease in performance across other test sets suggests that the signal transformation itself was harmful to performance.

We hypothesized that the decreased performance in the non-pig species was due to excessive input data loss. For example, through quantile normalization, the macaque dataset was reduced by one third, from 32, 697 reproducible peaks to 20, 615. We therefore created an extended quantile normalization (EQN) method that preserves the rank structure within each sample while aligning the overall signal distributions across species and accounts for different species having different numbers of peaks (**Materials and methods**). Unfortunately, models trained to predict EQN signals had comparable performance to those trained to predict log-transformed signals. Both models worked well on mouse chromosomes not used in training or hyper-parameter tuning (**Supplementary Table S8**) but had sub-optimal performance in non-mouse species.

When comparing the models trained to predict quantile-normalized or EQN signals, we observed that the EQN model strongly outperformed the quantile-normalized signal model on the macaque test sets. *r* increased from 0.23 to 0.35 and *ρ* increased from 0.23 to 0.37. The one exception was that the quantile-normalized signal model tended to predict that negatives had signals closer to 0 than the EQN model, though both models out-performed the models trained on log-transformed signals for this task.

Models trained to predict quantile-normalized, EQN, and log-transformed signals had comparably poor performance on their ability to predict differences in signal between species (**Supplementary Table S9**). While models trained on log-transformed signal consistently had lower mean squared error values, no model consistently outperformed the others across all four species, indicating that no single approach is superior for predicting chromatin accessibility across different species. These results demonstrate that normalizing signal across species is not sufficient for enabling models to accurately predict differences in chromatin accessibility between species.

### Training models with longer sequences does not improve performance

Large chromatin accessibility differences may be caused by sequence differences near peak summits, while smaller differences may be caused by sequence differences in flanking regions. We therefore trained a model using 2, 000 bp sequences centered on peak summits, quadrupling the size of our input sequences. Unfortunately, the new models showed no improvement over the original 500 bp models in their ability to predict signal strength or cross-species signal differences (**Supplementary Tables S8-S10**). In fact, the model trained using 2000 bp sequences made higher predictions for negatives than the models trained using 500 bp sequences. This suggests that longer input sequences do not enable models to make more accurate predictions of chromatin accessibility differences between species.

### Adding more species in training moderately improves performance

We and others previously found that liver chromatin accessibility classification models generalized to species not used in training and suggested that these results were indicative of a shared regulatory code across species (13, 14). However, in light of our new results, we hypothesized that the poor generalizability of regression models to new species may indicate that the regulatory code is only partially shared. Large differences in chromatin accessibility could reflect a shared regulatory code, while small differences may arise from species-specific regulatory elements. Alternatively, a previous study found that adding an additional species to model training for chromatin accessibility prediction improved performance on the original species, suggesting that predicting quantitative chromatin accessibility may require more training examples than predicting binary accessibility even if the regulatory code is shared (19).

To evaluate the extent to which incorporating more species into model training improves performance, we compared the performance of our model trained only on mouse data (“mouse-only model”) to a model that was also trained on data from macaque and rat (“3-species model”) as well as a model that was trained on data from all five species (“5-species model”). This allowed us to both evaluate the benefit from including more examples in training (evaluations on cow and pig for mouse-only versus 3-species model) as well as the necessity of including the evaluation species in training for achieving strong performance. We evaluated these models using the criteria described previously.

When evaluating predictions for the negative sets (**Supplementary Table S11**), the 5-species model was the most accurate at predicting values close to zero, with the 3-species model performing nearly as well. We observed a larger performance gap between the 3-species and the mouse-only models. Predictions for both positive and negative sets exhibited a negative shift for the 5-species model when compared to the 3-species model. However, the improvement from the mouse-only to 3-species model is not accompanied by a similar shift.

We observed a notable improvement in predicted and real signal correlations as well as MSE when comparing the mouse-only model to the 3-species model (**Supplementary Table S12**) across all species. The 3-species model exhibited a wider dynamic range and a line-of-best-fit slope more closely aligned with the identity line (y = x). In fact, *r* and *ρ* increased by approximately 0.1 on the mouse test set (**Fig. 4**). This suggests that training with more data causes the model to predict a wider range of values rather than clustering them around the mean. However, the improvement for the mouse model was smaller when increasing from 3 to 5 training species. The 3-species model performance for rat and macaque test sets were much better than the performance of the mouse-only model with *r* and *ρ* increasing by about 0.15 and 0.09 for macaque and rat, respectively (**Supplementary Fig. 3** and **Supplementary Fig. 4**). Interestingly, performance on macaque and rat slightly decreased when expanding to the 5-species model; while the *r* and *ρ* values were comparable, the 5-species model had a larger MSE and a smaller slope.

**Figure 4.**
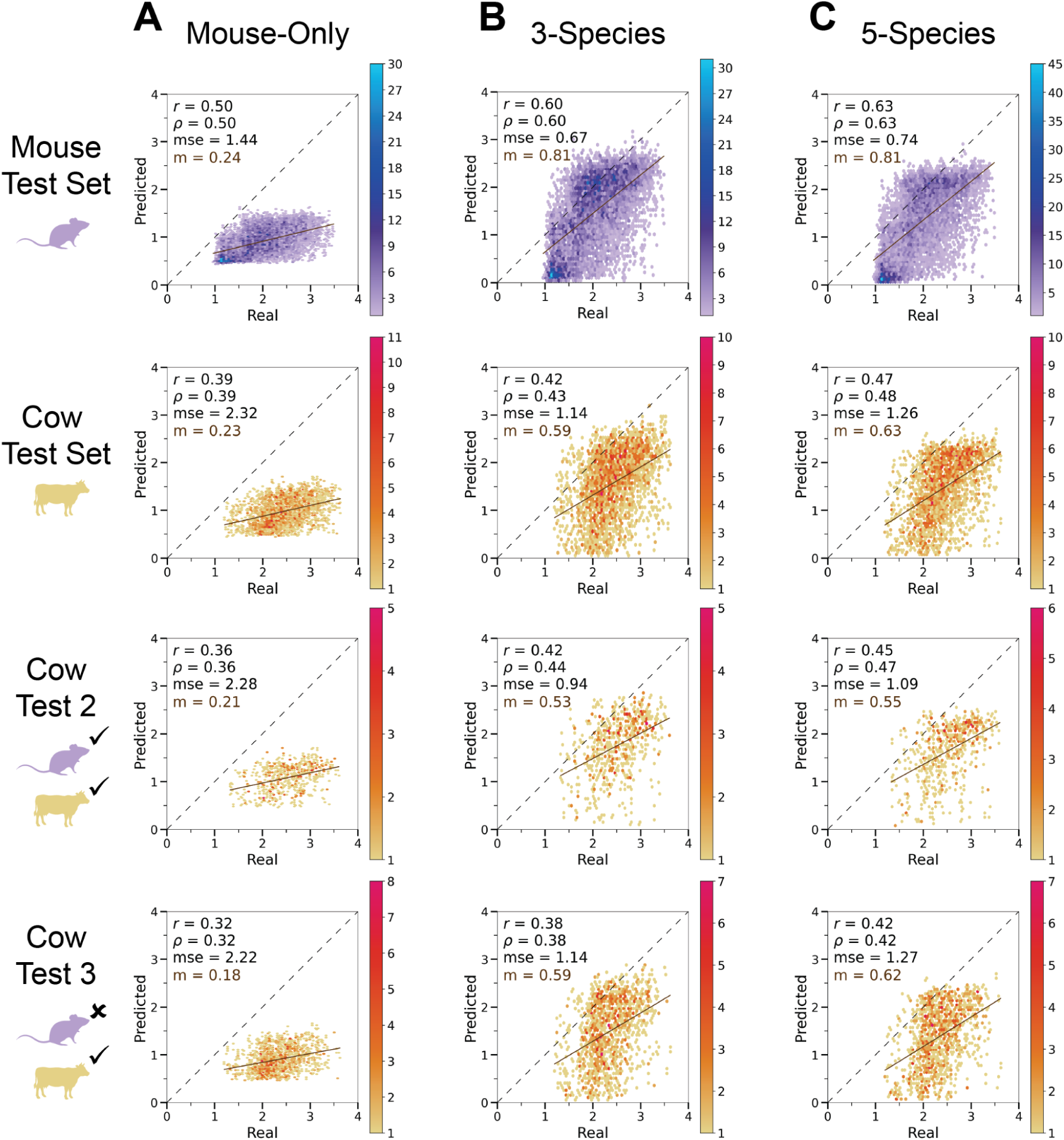
Binned scatter plots comparing real versus predicted signal values for positive examples of mouse and cow. The number of points in each bin is represented by the color scale. The dotted line represents perfect performance where points would align. The solid line indicates the line of best fit for the model’s predictions. (A) Prediction accuracy for mouse-only trained model. (B) for 3-species model (trained on mouse, macaque, and rat) (C) for 5-species model (trained on mouse, macaque, rat, cow, and pig). Metrics reported are the Pearson correlation coefficient (*r*), Spearman correlation coefficient (*ρ*), mean squared error (mse), and the slope of the line of best fit (*m*)

Performance of the 3-species model on the cow test sets was better than that of the mouse-only model (**Fig. 4**), while the performance difference for pig was smaller (**Supplementary Fig. 5**). Including cow data in the 5-species training set did not lead to a large performance increase on the cow test sets. For the “laurasiatheria-specific” set, we observed that the three-species model had better validation set but worse test set performance than the mouse-only model and that the 5-species model had only slightly better performance than the other models.

In addition, the 3-species model failed to predict chromatin accessibility differences between species, including between pairs of species used in training (**Supplementary Table S13**), and the performance of the 5-species model was generally not better for this task (**Fig. 3**). While all three models had comparable performance on Pearson and Spearman correlations, the mouse-only model had the lowest MSE. This demonstrates that including more training data, including data from the evaluation species, is insufficient for enabling models to predict differences in accessibility between species.

### Regulatory code differences between peaks with different levels of chromatin accessibility are often limited

We hypothesized that our models’ consistent poor performance on predicting accessibility differences between species may be a result of limited differences in regulatory code between peaks with different levels of chromatin accessibility. We performed standard and differential motif enrichment analysis on mouse candidate enhancers of strong, moderate, and weak strengths (**Materials and methods**). The differential motif enrichment between strong and weak candidate enhancers yielded some motifs, including motifs of TFs known to play important roles in the liver like HNF4A (56, 57) and CEBPA (56, 58) (**Data Availability**). However, the differential motif enrichment between strong and moderate candidate enhancers yielded limited results and only one motif of a TF that is known to be involved in liver-specific functions.

Looking at the individual motif enrichments, we found that strong and medium candidate enhancers had comparable enrichment for motifs of TFs with established roles in the liver. In contrast, weak candidate enhancers had much fewer enriched motifs; there were none for TFs with known roles in liver-specific functions, and there was no enrichment for CTCF, a TF integral to the chromatin looping required for enhancer function (59) . Given that the primary difference in motif composition is between strong/medium and weak regions, this may explain our model’s difficulty in predicting fine-grained quantitative differences. In other words, lack of clear sequence patterns that distinguish moderate from strong accessibility could be why the regression model struggles to predict these subtle signal differences between orthologous regions when both are accessible.

## DISCUSSION

Predicting differences in chromatin accessibility between orthologous regions across species is a critical yet previously underexplored task. To our knowledge, no prior work has evaluated regression models for their ability to predict differences in accessibility between orthologous regions. This task is necessary for using such models to connect chromatin accessibility differences between species when orthologs are accessible to phenotypic differences (44).

We hypothesized that training regression models to use DNA sequence to predict chromatin accessibility signal strength would enable us to predict differences in chromatin accessibility between species. Our models consistently had poor performance on our cross-species evaluation criteria, especially in predicting chromatin accessibility differences at orthologous regions, despite decent performance on parts of the training species not used in training. Across all models, including those trained with multiple species, performance for predicting accessibility differences between species remained consistent and suboptimal: models correctly predicted the direction of chromatin accessibility differences in just under 60% of orthologous sequences, and the correlation between predicted and real differences was under 0.40. We found that multiple sets of hyper-parameters and multiple input sequence lengths led to models with comparable performance. We also found that explicitly normalizing log-transformed chromatin accessibility signals across species did not yield better performance than using log-transformed signal values. We did, however, find that performance increased when predicting the EQN signal values when compared to quantile normalized values in macaque. Given that the quantile normalization procedure discards a substantial portion (approximately one third) of the macaque dataset, we hypothesize that this performance increase is a direct result of the EQN method’s ability to use the complete dataset. While performance was consistently above random chance, it remains too low to be reliably used in applications such as linking chromatin accessibility differences to phenotypic differences between species (44).

Our results additionally demonstrated that incorporating data from more species into model training generally improves model performance for both species used in training and for held-out species. As a previous study suggested (19), we hypothesize that this occurs because the liver regulatory code is shared across species, thus adding data from additional species exposes the models to more diverse yet relevant training examples. However, we found the improvement from incorporating additional species into model training was limited when comparing the 3-species to the 5-species model. One possibility is that having three species of training data is sufficient for training a model with the best possible performance. Alternatively, the lower quality of the cow and pig data added to model training may be unable to help improve model performance. In addition, the performance on ability to predict differences in chromatin accessibility between species was poor regardless of the number of species used when training the model.

While our regression models can usually correctly identify negative sequences, performance on positive sequences, particularly in terms of accuracy of predicted accessibility signal strength in species not used in training and predicted differences between species in accessibility, remains weak. The limited improvement in predicting quantitative chromatin accessibility differences from extensive model modifications, including hyperparameter tuning, alternative signal normalization, longer input sequences, and multi-species training suggests that the task may be infeasible. This infeasibility may be a result of small differences between liver cell type proportions or differences between percentages of hepatocytes in each liver lobule zone between species (60). Alternatively, poor performance may reflect other differences between samples from different species, such as slight variations in age (all were adults) or the time of sacrifice within their circadian cycles. However, the lack of motif enrichment differences between strong and moderate peaks within a species suggests that training a model to distinguish them by sequence alone may be infeasible, even if cell type proportions were identical and samples were matched more closely by age, circadian timing, and other factors across species. This lack of motif enrichment in contrast to the liver motif enrichment between strong and weak peaks may explain why our regression models were unsuccessful despite the success of classification models we previously trained (13, 44).

There are multiple reasons why few sequence patterns differentiate peaks with moderate versus strong levels of accessibility. One is that these differences may not be driven by DNA sequence. Specific combinations of sequence motifs may be required for accessibility in a subset of cells, while broader accessibility could depend on tissue environments not encoded in the genome. Another possibility is that proper quantification of differences between moderate and strong accessibility requires deeper sequencing than we had from many of our animals or a different approach to quantifying signals in peaks than the one used by MACS2. Finally, these differences might be influenced by regulatory sequences located more than 1 kb from the peak summit—such as flanking regions or distal elements that interact with enhancers through three-dimensional chromatin looping. The upcoming assembly and alignment of hundreds of chromosome-level vertebrate genome assemblies by the Vertebrate Genomes Project (22) and Bat1K Project (23) will provide an opportunity for us to evaluate the ability of models trained using much longer sequences to predict differences between species in chromatin accessibility and mitigate errors caused by sub-optimal genome assembly quality (21). Unfortunately, most three-dimensional interaction data from primary tissues lacks the resolution to identify enhancer interactions that may affect chromatin accessibility (61). For example, to our knowledge, no Micro-C data exists for the liver. If generated, incorporating such data into model training could be a valuable extension of this work.

An additional unexpected result was that our mouse-only trained models consistently had the worst performance in the rat, including when evaluating the ability to predict the accessibility signal of peaks in the rat whose orthologs are accessible in the mouse. This was especially surprising given that the rat is more closely related to the mouse than the other species are. One explanation is that the second rat replicate was extremely shallow (**Supplementary Table S1**) and the first was not especially deep, making the accessibility quantification potentially unreliable. Other possibilities include that the cell type proportions sampled from the rats might have been different from those sampled for other species or that differences between the way that the rats and the other animals were raised affected the rats’ chromatin accessibility.

Our evaluation criteria were specifically designed to assess a model’s ability to predict accessibility at orthologs of accessible sites. Evaluating genome-wide chromatin accessibility predictions in a given tissue would require additional criteria, such as verifying that the model correctly predicts that regions inaccessible in the target tissue (but accessible in others) are indeed inaccessible. While many previous classification models have shown strong performance on held-out chromosomes, only some have performed well on this task (13). Additionally, genome-wide prediction would require a strategy for defining input sequences. Our models used sequences centered on peak summit orthologs, but such anchors do not exist for regions that are not orthologs of accessible sites.

This work highlights the importance of evaluating machine learning models for genomics using criteria that are directly applicable to how they will be used. We found that models that provide accurate quantitative predictions of chromatin accessibility on held-out chromosomes consistently perform better than random but poorly in predicting chromatin accessibility differences between species. Being able to predict these differences is necessary for using predictions to study differences between species.

These results are in line with our previous work that demonstrated that only a strict subset of models that can predict presence of an accessible region in held-out chromosomes can predict difference in accessibility status between orthologous regions (13). Work from others reveal that models that accurately predict gene expression for held-out chromosomes fail to predict the effects of genetic differences between people in gene expression (26, 27, 62), further demonstrating the importance of directly evaluating machine learning models’ abilities to predict the effects of changes in sequence. However, the lack of sequence patterns enriched for occurring in strong peaks relative to moderate peaks suggest that there might be limited biological value in obtaining precise predictions of chromatin accessibility level above a certain value, as small and even moderate differences in chromatin accessibility between species may have little impact on phenotype evolution.

## ACKNOWLEDGEMENTS

We are thankful to all present and past members of the Pfenning Kaplow Labs for useful discussions. We are grateful to J. He for his assistance with the macaque sample collection. This work used the Extreme Science and Engineering Discovery Environment (XSEDE), through the Pittsburgh Supercomputing Center Bridges and Bridges-2 Compute Clusters, which was supported by National Science Foundation grants TG-BIO200055 and ACI-1548562 (63).

## Author contributions

I.M.K. and A.R.P. conceptualized the project and provided overall supervision. A.S. performed the computational analyses and prepared the manuscript with assistance from A.R.B. and I.M.K. H.H.S. and I.M.K. supervised software development. A.R. contributed to the initial project framework. M.E.W., A.J.L., A.R.B., and W.R.S. conducted the animal experiments. All authors reviewed and approved the final manuscript.

## DATA AVAILABILITY

Macaque and rat ATAC-seq fastq, bigWig, and narrowPeak have been made available at GEO under accession ID GSE159815. This manuscript also used publicly available data obtained from GSM4847918, GSM4847919, and GSE158430.

Full quality control evaluations and motif discovery results can be found at http://daphne.compbio.cs.cmu.edu/files/azstephe/liver_regression_resource/.

