## Supplementary Information for "Challenges in Predicting Chromatin Accessibility Differences between Species"

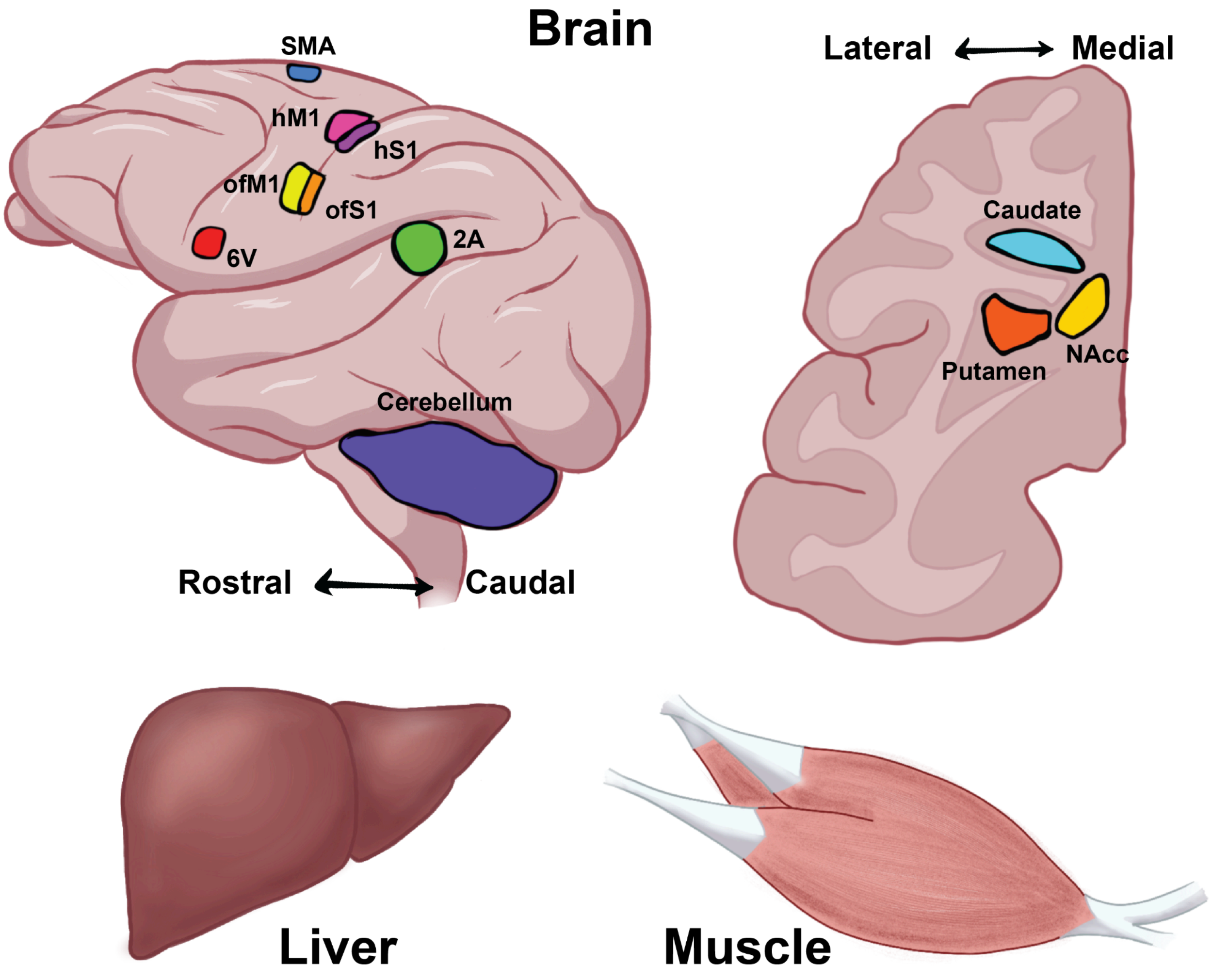

**Figure S1.** Diagrams show the liver and specific brain regions sampled from rhesus macaque and rat for this study. Brain regions sampled from macaque include secondary auditory cortex (2A), premotor area 6V (6V), supplementary motor area (SMA), hand and forearm motor cortex (hM1), hand and forearm S1 (hS1), orofacial motor cortex (ofM1), orofacial S1 (ofS1), cerebellum, caudate, nucleus accumbens (NAcc), and putamen. Samples collected from rat include liver, primary motor cortex (M1), and caudoputamen of the striatum (STR).

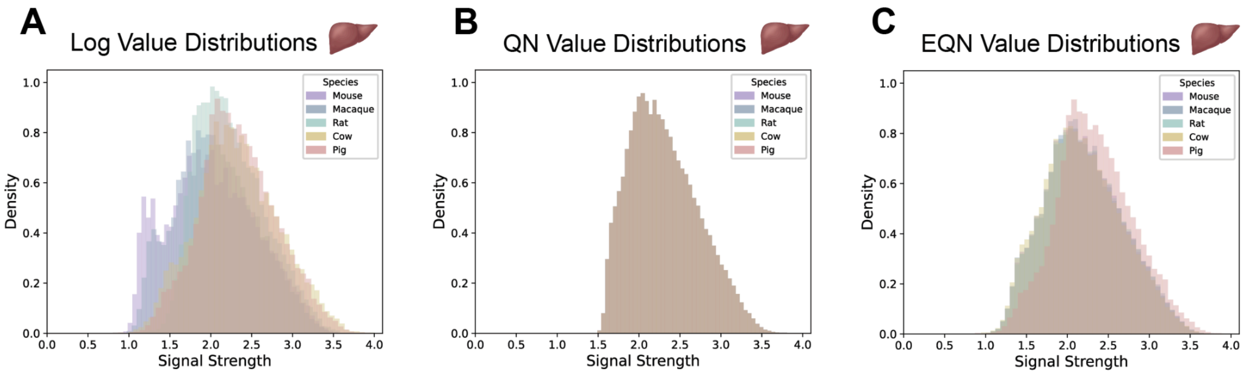

**Figure S2.** Histograms illustrate the distribution of observed signal values after three preprocessing methods for the five species: **(A)** log-transformation, **(B)** Quantile Normalization (QN), and **(C)** Extended Quantile Normalization (EQN).

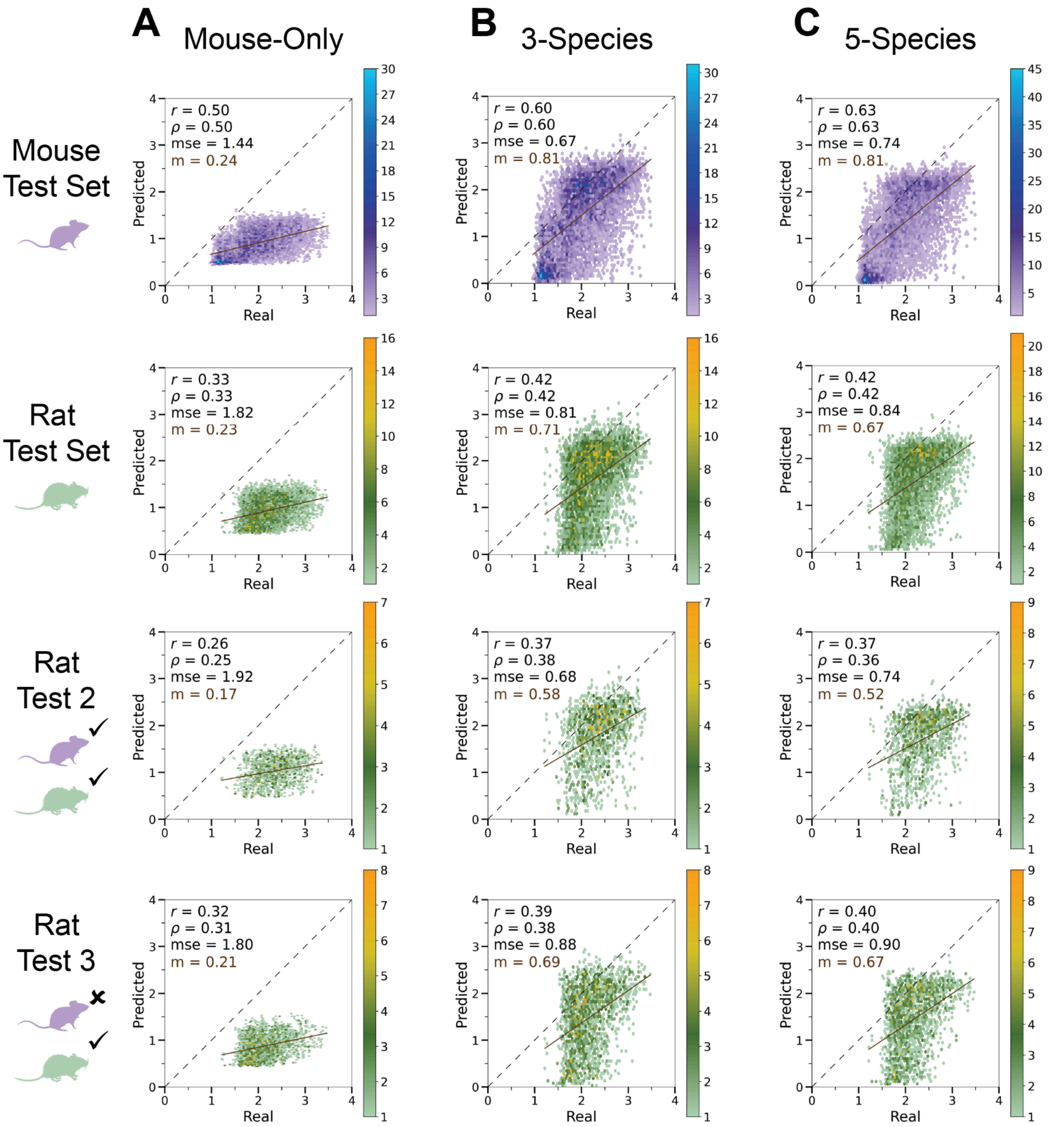

**Figure S3.** Binned scatter plots comparing real versus predicted signal values for positive examples of mouse and rat. The number of points in each bin is represented by the color scale. The dotted line represents perfect performance where points would align. The solid line indicates the line of best fit for the model's predictions. (A) Prediction accuracy for mouse-only trained model. (B) for 3-species model (trained on mouse, macaque, and rat) (C) for 5-species model (trained on mouse, macaque, rat, cow, and pig). Metrics reported are the Pearson correlation coefficient (*r*), Spearman correlation coefficient (*ρ*), mean squared error (mse), and the slope of the line of best fit (*m*).

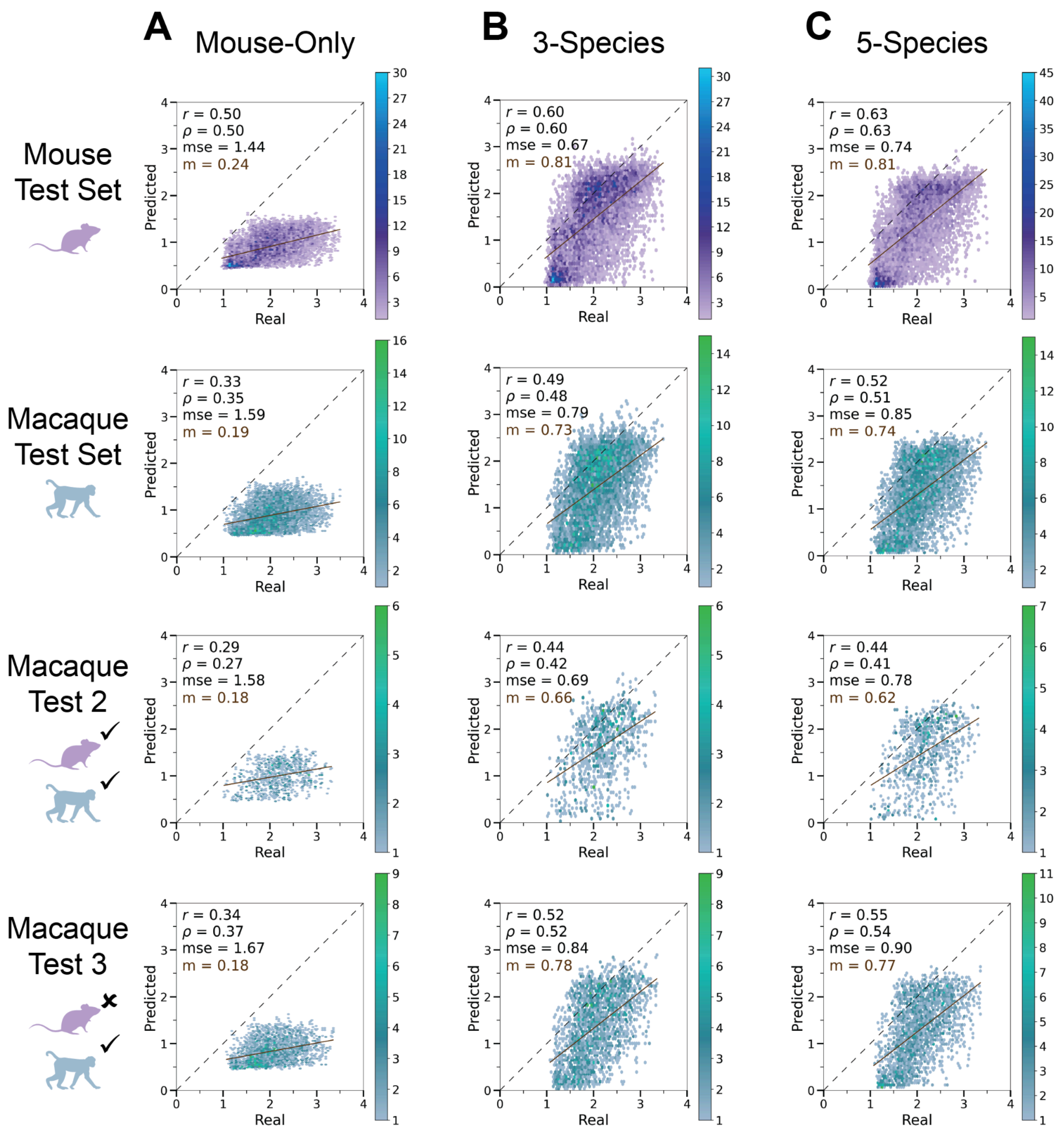

**Figure S4.** Binned scatter plots comparing real versus predicted signal values for positive examples of mouse and macaque. The number of points in each bin is represented by the color scale. The dotted line represents perfect performance where points would align. The solid line indicates the line of best fit for the model's predictions. (A) Prediction accuracy for mouse-only trained model. (B) for 3-species model (trained on mouse, macaque, and rat) (C) for 5-species model (trained on mouse, macaque, rat, cow, and pig). Metrics reported are the Pearson correlation coefficient ($r$), Spearman correlation coefficient ($\rho$), mean squared error (mse), and the slope of the line of best fit ($m$).

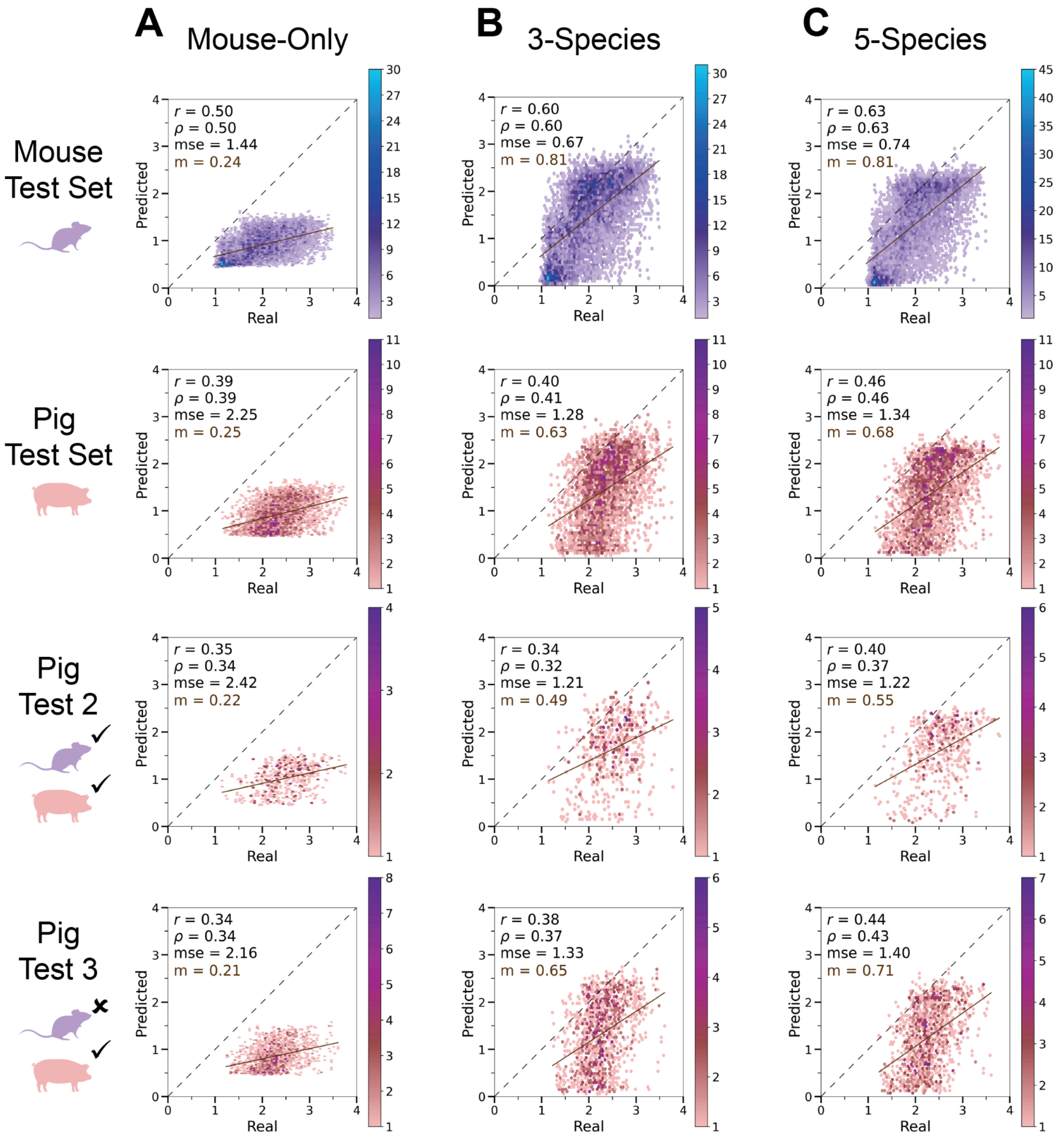

**Figure S5.** Binned scatter plots comparing real versus predicted signal values for positive examples of mouse and pig. The number of points in each bin is represented by the color scale. The dotted line represents perfect performance where points would align. The solid line indicates the line of best fit for the model's predictions. (A) Prediction accuracy for mouse-only trained model. (B) for 3-species model (trained on mouse, macaque, and rat) (C) for 5-species model (trained on mouse, macaque, rat, cow, and pig). Metrics reported are the Pearson correlation coefficient (*r*), Spearman correlation coefficient (*ρ*), mean squared error (mse), and the slope of the line of best fit (*m*).

**Table S1.**

| Species | Dataset Source | Number of Distinct Fragments, Rep. 1 | Number of Distinct Fragments, Rep. 2 |
|---|---|---|---|
| *Mus musculus* | (13) | 21,913,403 | 38,064,998 |
| *Rattus norvegicus* | This paper | 33,735,211 | 11,175,366 |
| *Macaca mulatta* | This paper | 104,947,535 | Not used due to poor data quality |
| *Bos taurus* | (29) | 19,716,631 | 18,350,665 |
| *Sus scrofa* | (29) | 15,823,907 | 24,024,007 |

**Table S2.** Size of positive and negative sets across training, validation, and test splits by species

| Set | Total | Training | Validation | Test |
|---|---|---|---|---|
| Mouse Positive | 22042 | 16250 | 2016 | 3776 |
| Mouse Negative | | 18851 | 2587 | 3698 |
| Macaque Positive | 32697 | 12067 | 1788 | 2624 |
| Macaque Negative | | 16165 | 1943 | 3479 |
| Rat Positive | 21531 | 13365 | 1533 | 2576 |
| Rat Negative | | 19408 | 2850 | 4405 |
| Cow Positive | 25806 | 7687 | 1162 | 1664 |
| Cow Negative | | 17403 | 2049 | 3720 |
| Pig Positive | 20615 | 7762 | 992 | 1627 |
| Pig Negative | | 14436 | 1770 | 3061 |

**Table S3.** Hyperparameters Comparison of the 5 best mouse-only models

| Hyperparameters | Best Mouse-Only | Alternate 1 | Alternate 2 | Alternate 3 | Alternate 4 |
|---|---|---|---|---|---|
| Conv Filters | 200 | 175 | 175 | 300 | 225 |
| Dense Filters | 250 | 200 | 100 | 150 | 100 |
| Conv Dropout | 0.55 | 0.55 | 0.55 | 0.5 | 0.6 |
| Dense Dropout | 0.2 | 0.4 | 0.5 | 0.45 | 0.2 |
| Conv L2 | 1E-0.4 | 1E-0.4 | 1E-0.4 | 1E-0.4 | 1E-0.4 |
| Dense L2 | 1E-0.4 | 1E-0.4 | 1E-0.4 | 1E-0.4 | 1E-0.4 |
| Initial LR | 6.14E-04 | 8.27E-04 | 5.50E-04 | 1.42E-03 | 7.26E-04 |
| Conv Layers | 6 | 2 | 2 | 2 | 2 |
| Dense Layers | 2 | 1 | 1 | 1 | 1 |

Conv Filters: Number of filters per convolutional layer; Conv Dropout: Convolutional layer dropout; Dense Dropout: Fully connected layer dropout; Conv L2: Convolutional layer L2 regularization; Dense L2: Fully connected layer L2 regularization; Initial LR: Initial learning rate, Conv Layers: Number of convolutional layers; Dense Layers: Number of fully connected layers.

**Table S4.** Hyperparameter comparison of the best models

| Hyperparameters | Best Mouse-Only | QN | EQN | 2000 bp | 3-Species | 5-Species |
|---|---|---|---|---|---|---|
| Conv Filters | 200 | 300 | 375 | 275 | 225 | 275 |
| Dense Filters | 250 | 150 | 200 | 300 | 250 | 350 |
| Conv Dropout | 0.55 | 0.3 | 0.4 | 0.55 | 0.2 | 0.25 |
| Dense Dropout | 0.2 | 0.4 | 0.4 | 0.4 | 0.01 | 0.35 |
| Conv L2 | 1E-0.4 | 1E-0.4 | 1E-0.4 | 1E-0.4 | 2.64E-05 | 4.05E-04 |
| Dense L2 | 1E-0.4 | 1E-0.4 | 1E-0.4 | 1E-0.4 | 2.41E-04 | 1.28E-03 |
| Initial LR | 6.14E-04 | 2.79E-0.4 | 1.18E-04 | 6.35E-04 | 1.17E-04 | 3.49E-05 |
| Conv Layers | 6 | 5 | 2 | 3 | 4 | 5 |
| Dense Layers | 2 | 1 | 2 | 1 | 3 | 4 |

Conv Filters: Number of filters per convolutional layer; Conv Dropout: Convolutional layer dropout; Dense Dropout: Fully connected layer dropout; Conv L2: Convolutional layer L2 regularization; Dense L2: Fully connected layer L2 regularization; Initial LR: Initial learning rate, Conv Layers: Number of convolutional layers; Dense Layers: Number of fully connected layers.

**Table S5.** Performance of the 5 best mouse-only trained models

| Species | Group | Metric | Best Mouse-Only | Alternate 1 | Alternate 2 | Alternate 3 | Alternate 4 | Number of Peaks |
|---|---|---|---|---|---|---|---|---|
| Mouse | Train | Pearson | 0.494 | 0.497 | 0.507 | 0.485 | 0.478 | 16250 |
| Mouse | Train | Pearson P-Val | 0.00E+00 | 0.00E+00 | 0.00E+00 | 0.00E+00 | 0.00E+00 | |
| Mouse | Train | Spearman | 0.498 | 0.509 | 0.521 | 0.502 | 0.501 | 16250 |
| Mouse | Train | Spearman P-Val | 0.00E+00 | 0.00E+00 | 0.00E+00 | 0.00E+00 | 0.00E+00 | |
| Mouse | Train | MSE | 1.39 | 1.07 | 1.02 | 1.07 | 1.18 | |
| Mouse | Validation | Pearson | 0.483 | 0.457 | 0.465 | 0.458 | 0.451 | 2016 |
| Mouse | Validation | Pearson P-Val | 3.49E-232 | 2.26E-205 | 3.68E-213 | 4.93E-206 | 1.25E-198 | |
| Mouse | Validation | Spearman | 0.491 | 0.477 | 0.488 | 0.48 | 0.484 | 2016 |
| Mouse | Validation | Spearman P-Val | 8.59E-242 | 1.10E-225 | 4.58E-238 | 2.25E-229 | 2.39E-233 | |
| Mouse | Validation | MSE | 1.42 | 1.13 | 1.07 | 1.11 | 1.22 | |
| Mouse | Test | Pearson | 0.496 | 0.48 | 0.496 | 0.48 | 0.483 | 3776 |
| Mouse | Test | Pearson P-Val | 0.00E+00 | 0.00E+00 | 0.00E+00 | 0.00E+00 | 0.00E+00 | |
| Mouse | Test | Spearman | 0.502 | 0.49 | 0.509 | 0.498 | 0.504 | 3776 |
| Mouse | Test | Spearman P-Val | 0.00E+00 | 0.00E+00 | 0.00E+00 | 0.00E+00 | 0.00E+00 | |
| Mouse | Test | MSE | 1.44 | 1.15 | 1.09 | 1.13 | 1.23 | |
| Cow | Train | Pearson | 0.406 | 0.39 | 0.387 | 0.384 | 0.388 | 7687 |
| Cow | Train | Pearson P-Val | 0.00E+00 | 0.00E+00 | 0.00E+00 | 0.00E+00 | 0.00E+00 | |
| Cow | Train | Spearman | 0.401 | 0.385 | 0.381 | 0.375 | 0.381 | 7687 |
| Cow | Train | Spearman P-Val | 0.00E+00 | 0.00E+00 | 0.00E+00 | 0.00E+00 | 0.00E+00 | |
| Cow | Train | MSE | 2.27 | 1.87 | 1.81 | 1.79 | 1.94 | |
| Cow | Validation | Pearson | 0.4 | 0.407 | 0.397 | 0.386 | 0.393 | 1162 |
| Cow | Validation | Pearson P-Val | 7.22E-88 | 3.67E-91 | 1.81E-86 | 2.49E-81 | 2.27E-84 | |
| Cow | Validation | Spearman | 0.39 | 0.399 | 0.39 | 0.377 | 0.387 | 1162 |
| Cow | Validation | Spearman P-Val | 6.12E-83 | 1.81E-87 | 7.80E-83 | 5.46E-77 | 8.93E-82 | |
| Cow | Validation | MSE | 2.15 | 1.79 | 1.72 | 1.7 | 1.83 | |

| | | | | | | | | |
|---|---|---|---|---|---|---|---|---|
| Cow | Test | Pearson | 0.385 | 0.367 | 0.362 | 0.358 | 0.368 | 1284 |
| Cow | Test | Pearson P-Val | 2.67E-89 | 2.41E-80 | 2.85E-78 | 2.07E-76 | 4.90E-81 | |
| Cow | Test | Spearman | 0.391 | 0.375 | 0.368 | 0.365 | 0.379 | 1284 |
| Cow | Test | Spearman P-Val | 1.77E-92 | 2.83E-84 | 5.89E-81 | 1.53E-79 | 3.93E-86 | |
| Cow | Test | MSE | 2.32 | 1.96 | 1.89 | 1.85 | 2.01 | |
| Cow | Val2 | Pearson | 0.547 | 0.511 | 0.489 | 0.478 | 0.49 | 194 |
| Cow | Val2 | Pearson P-Val | 2.47E-29 | 7.13E-25 | 2.14E-22 | 2.83E-21 | 1.64E-22 | |
| Cow | Val2 | Spearman | 0.54 | 0.504 | 0.473 | 0.475 | 0.482 | 194 |
| Cow | Val2 | Spearman P-Val | 2.17E-28 | 4.04E-24 | 9.06E-21 | 5.87E-21 | 1.10E-21 | |
| Cow | Val2 | MSE | 2.26 | 1.74 | 1.73 | 1.7 | 1.87 | |
| Cow | Val3 | Pearson | 0.256 | 0.267 | 0.279 | 0.272 | 0.278 | 556 |
| Cow | Val3 | Pearson P-Val | 8.24E-16 | 2.56E-17 | 4.42E-19 | 4.82E-18 | 6.84E-19 | |
| Cow | Val3 | Spearman | 0.263 | 0.288 | 0.298 | 0.283 | 0.301 | 556 |
| Cow | Val3 | Spearman P-Val | 8.16E-17 | 2.35E-20 | 5.99E-22 | 1.10E-19 | 2.17E-22 | |
| Cow | Val3 | MSE | 2.12 | 1.87 | 1.73 | 1.73 | 1.84 | |
| Cow | Test2 | Pearson | 0.364 | 0.355 | 0.343 | 0.357 | 0.355 | 337 |
| Cow | Test2 | Pearson P-Val | 3.37E-20 | 4.05E-19 | 8.55E-18 | 1.89E-19 | 4.04E-19 | |
| Cow | Test2 | Spearman | 0.364 | 0.351 | 0.341 | 0.352 | 0.36 | 337 |
| Cow | Test2 | Spearman P-Val | 3.46E-20 | 1.26E-18 | 1.73E-17 | 9.78E-19 | 9.99E-20 | |
| Cow | Test2 | MSE | 2.28 | 1.79 | 1.8 | 1.78 | 1.91 | |
| Cow | Test3 | Pearson | 0.319 | 0.303 | 0.324 | 0.312 | 0.315 | 694 |
| Cow | Test3 | Pearson P-Val | 8.43E-32 | 1.27E-28 | 4.46E-33 | 2.59E-30 | 4.12E-31 | |
| Cow | Test3 | Spearman | 0.321 | 0.304 | 0.318 | 0.305 | 0.304 | 694 |
| Cow | Test3 | Spearman P-Val | 2.20E-32 | 9.15E-29 | 9.77E-32 | 6.38E-29 | 7.61E-29 | |
| Cow | Test3 | MSE | 2.22 | 1.95 | 1.81 | 1.79 | 1.94 | |
| Macaque | Train | Pearson | 0.361 | 0.365 | 0.378 | 0.358 | 0.352 | 12066 |
| Macaque | Train | Pearson P-Val | 0.00E+00 | 0.00E+00 | 0.00E+00 | 0.00E+00 | 0.00E+00 | |

| | | | | | | | | |
|---|---|---|---|---|---|---|---|---|
| Macaque | Train | Spearman | 0.374 | 0.383 | 0.394 | 0.376 | 0.376 | 12066 |
| Macaque | Train | Spearman P-Val | 0.00E+00 | 0.00E+00 | 0.00E+00 | 0.00E+00 | 0.00E+00 | |
| Macaque | Train | MSE | 1.6 | 1.3 | 1.22 | 1.24 | 1.34 | |
| Macaque | Validation | Pearson | 0.355 | 0.366 | 0.379 | 0.36 | 0.354 | 1788 |
| Macaque | Validation | Pearson P-Val | 3.57E-104 | 1.40E-111 | 2.65E-120 | 2.69E-107 | 1.01E-103 | |
| Macaque | Validation | Spearman | 0.366 | 0.387 | 0.398 | 0.381 | 0.382 | 1788 |
| Macaque | Validation | Spearman P-Val | 1.46E-111 | 9.37E-126 | 1.05E-133 | 2.80E-121 | 2.37E-122 | |
| Macaque | Validation | MSE | 1.56 | 1.27 | 1.19 | 1.21 | 1.3 | |
| Macaque | Test | Pearson | 0.327 | 0.339 | 0.36 | 0.339 | 0.325 | 2624 |
| Macaque | Test | Pearson P-Val | 2.07E-128 | 1.28E-138 | 4.04E-158 | 6.95E-139 | 1.06E-126 | |
| Macaque | Test | Spearman | 0.346 | 0.365 | 0.383 | 0.364 | 0.355 | 2624 |
| Macaque | Test | Spearman P-Val | 2.89E-145 | 5.65E-163 | 7.32E-181 | 1.79E-161 | 3.90E-153 | |
| Macaque | Test | MSE | 1.59 | 1.31 | 1.23 | 1.25 | 1.35 | |
| Macaque | Val2 | Pearson | 0.437 | 0.399 | 0.413 | 0.393 | 0.38 | 331 |
| Macaque | Val2 | Pearson P-Val | 5.34E-30 | 2.54E-24 | 2.00E-26 | 1.65E-23 | 7.68E-22 | |
| Macaque | Val2 | Spearman | 0.431 | 0.423 | 0.449 | 0.418 | 0.424 | 331 |
| Macaque | Val2 | Spearman P-Val | 4.72E-29 | 7.77E-28 | 8.50E-32 | 3.96E-27 | 6.11E-28 | |
| Macaque | Val2 | MSE | 1.55 | 1.21 | 1.18 | 1.2 | 1.31 | |
| Macaque | Val3 | Pearson | 0.33 | 0.378 | 0.382 | 0.368 | 0.358 | 887 |
| Macaque | Val3 | Pearson P-Val | 5.18E-44 | 4.88E-59 | 2.96E-60 | 9.68E-56 | 1.39E-52 | |
| Macaque | Val3 | Spearman | 0.345 | 0.39 | 0.392 | 0.386 | 0.378 | 887 |
| Macaque | Val3 | Spearman P-Val | 1.49E-48 | 3.76E-63 | 6.74E-64 | 7.73E-62 | 4.54E-59 | |
| Macaque | Val3 | MSE | 1.64 | 1.36 | 1.25 | 1.27 | 1.37 | |
| Macaque | Test2 | Pearson | 0.289 | 0.282 | 0.305 | 0.287 | 0.267 | 472 |
| Macaque | Test2 | Pearson P-Val | 2.23E-17 | 1.83E-16 | 1.65E-19 | 5.16E-17 | 1.33E-14 | |
| Macaque | Test2 | Spearman | 0.274 | 0.279 | 0.299 | 0.291 | 0.27 | 472 |
| Macaque | Test2 | Spearman P-Val | 2.24E-15 | 4.98E-16 | 1.37E-18 | 1.24E-17 | 6.18E-15 | |

| | | | | | | | | |
|---|---|---|---|---|---|---|---|---|
| Macaque | Test2 | MSE | 1.58 | 1.24 | 1.2 | 1.23 | 1.35 | |
| Macaque | Test3 | Pearson | 0.342 | 0.388 | 0.411 | 0.384 | 0.384 | 1263 |
| Macaque | Test3 | Pearson P-Val | 6.17E-68 | 4.70E-89 | 1.97E-101 | 3.15E-87 | 3.77E-87 | |
| Macaque | Test3 | Spearman | 0.369 | 0.411 | 0.431 | 0.396 | 0.403 | 1263 |
| Macaque | Test3 | Spearman P-Val | 5.09E-80 | 2.97E-101 | 2.14E-112 | 1.76E-93 | 9.44E-97 | |
| Macaque | Test3 | MSE | 1.67 | 1.4 | 1.28 | 1.32 | 1.41 | |
| Pig | Train | Pearson | 0.355 | 0.351 | 0.345 | 0.342 | 0.345 | 7762 |
| Pig | Train | Pearson P-Val | 0.00E+00 | 0.00E+00 | 0.00E+00 | 0.00E+00 | 0.00E+00 | |
| Pig | Train | Spearman | 0.347 | 0.347 | 0.34 | 0.336 | 0.342 | 7762 |
| Pig | Train | Spearman P-Val | 0.00E+00 | 0.00E+00 | 0.00E+00 | 0.00E+00 | 0.00E+00 | |
| Pig | Train | MSE | 2.26 | 1.9 | 1.83 | 1.81 | 1.94 | |
| Pig | Validation | Pearson | 0.394 | 0.372 | 0.362 | 0.35 | 0.349 | 992 |
| Pig | Validation | Pearson P-Val | 3.47E-72 | 5.48E-64 | 2.61E-60 | 6.59E-56 | 1.02E-55 | |
| Pig | Validation | Spearman | 0.394 | 0.38 | 0.367 | 0.357 | 0.355 | 992 |
| Pig | Validation | Spearman P-Val | 2.08E-72 | 6.02E-67 | 6.30E-62 | 2.29E-58 | 1.50E-57 | |
| Pig | Validation | MSE | 2.08 | 1.75 | 1.68 | 1.67 | 1.78 | |
| Pig | Test | Pearson | 0.388 | 0.367 | 0.354 | 0.355 | 0.361 | 1627 |
| Pig | Test | Pearson P-Val | 2.24E-115 | 8.76E-102 | 3.09E-94 | 9.65E-95 | 2.39E-98 | |
| Pig | Test | Spearman | 0.394 | 0.372 | 0.359 | 0.355 | 0.369 | 1627 |
| Pig | Test | Spearman P-Val | 2.51E-119 | 4.89E-105 | 4.77E-97 | 4.33E-95 | 3.80E-103 | |
| Pig | Test | MSE | 2.25 | 1.91 | 1.84 | 1.82 | 1.94 | |
| Pig | Val2 | Pearson | 0.456 | 0.381 | 0.387 | 0.385 | 0.369 | 164 |
| Pig | Val2 | Pearson P-Val | 6.06E-16 | 1.71E-10 | 7.54E-11 | 9.64E-11 | 1.08E-09 | |
| Pig | Val2 | Spearman | 0.45 | 0.377 | 0.376 | 0.382 | 0.365 | 164 |
| Pig | Val2 | Spearman P-Val | 1.79E-15 | 3.35E-10 | 3.49E-10 | 1.61E-10 | 1.84E-09 | |
| Pig | Val2 | MSE | 2.14 | 1.66 | 1.66 | 1.64 | 1.76 | |
| Pig | Val3 | Pearson | 0.274 | 0.304 | 0.288 | 0.281 | 0.28 | 475 |

| | | | | | | | | |
|---|---|---|---|---|---|---|---|---|
| Pig | Val3 | Pearson P-Val | 1.51E-15 | 1.92E-19 | 2.54E-17 | 2.06E-16 | 2.59E-16 | |
| Pig | Val3 | Spearman | 0.269 | 0.3 | 0.279 | 0.272 | 0.27 | 475 |
| Pig | Val3 | Spearman P-Val | 5.79E-15 | 7.37E-19 | 3.65E-16 | 3.07E-15 | 5.49E-15 | |
| Pig | Val3 | MSE | 2.08 | 1.82 | 1.71 | 1.69 | 1.79 | |
| Pig | Test2 | Pearson | 0.351 | 0.293 | 0.283 | 0.277 | 0.293 | 293 |
| Pig | Test2 | Pearson P-Val | 4.29E-16 | 9.78E-11 | 6.18E-10 | 1.76E-09 | 9.45E-11 | |
| Pig | Test2 | Spearman | 0.342 | 0.29 | 0.279 | 0.279 | 0.308 | 293 |
| Pig | Test2 | Spearman P-Val | 3.58E-15 | 1.59E-10 | 1.21E-09 | 1.33E-09 | 5.01E-12 | |
| Pig | Test2 | MSE | 2.42 | 1.96 | 1.96 | 1.94 | 2.06 | |
| Pig | Test3 | Pearson | 0.337 | 0.335 | 0.332 | 0.333 | 0.326 | 716 |
| Pig | Test3 | Pearson P-Val | 4.27E-37 | 1.23E-36 | 7.47E-36 | 5.08E-36 | 1.73E-34 | |
| Pig | Test3 | Spearman | 0.337 | 0.327 | 0.329 | 0.322 | 0.323 | 716 |
| Pig | Test3 | Spearman P-Val | 3.89E-37 | 8.31E-35 | 3.61E-35 | 1.59E-33 | 9.35E-34 | |
| Pig | Test3 | MSE | 2.16 | 1.91 | 1.8 | 1.77 | 1.88 | |
| Rat | Train | Pearson | 0.327 | 0.322 | 0.321 | 0.315 | 0.306 | 13365 |
| Rat | Train | Pearson P-Val | 0.00E+00 | 0.00E+00 | 0.00E+00 | 0.00E+00 | 0.00E+00 | |
| Rat | Train | Spearman | 0.331 | 0.33 | 0.327 | 0.322 | 0.317 | 13365 |
| Rat | Train | Spearman P-Val | 0.00E+00 | 0.00E+00 | 0.00E+00 | 0.00E+00 | 0.00E+00 | |
| Rat | Train | MSE | 1.89 | 1.53 | 1.46 | 1.49 | 1.61 | |
| Rat | Validation | Pearson | 0.301 | 0.284 | 0.287 | 0.276 | 0.269 | 1533 |
| Rat | Validation | Pearson P-Val | 9.10E-63 | 1.38E-55 | 7.17E-57 | 1.59E-52 | 1.04E-49 | |
| Rat | Validation | Spearman | 0.304 | 0.294 | 0.296 | 0.291 | 0.28 | 1533 |
| Rat | Validation | Spearman P-Val | 2.19E-64 | 8.56E-60 | 5.96E-61 | 1.73E-58 | 4.37E-54 | |
| Rat | Validation | MSE | 1.84 | 1.49 | 1.42 | 1.44 | 1.57 | |
| Rat | Test | Pearson | 0.333 | 0.32 | 0.321 | 0.307 | 0.311 | 2576 |
| Rat | Test | Pearson P-Val | 5.67E-131 | 2.23E-120 | 1.03E-121 | 1.72E-110 | 1.72E-113 | |
| Rat | Test | Spearman | 0.332 | 0.327 | 0.329 | 0.321 | 0.325 | 2576 |

| | | | | | | | | |
|---|---|---|---|---|---|---|---|---|
| Rat | Test | Spearman P-Val | 1.07E-130 | 9.60E-127 | 6.53E-128 | 3.75E-121 | 2.25E-124 | |
| Rat | Test | MSE | 1.82 | 1.5 | 1.43 | 1.46 | 1.57 | |
| Rat | Val2 | Pearson | 0.226 | 0.196 | 0.18 | 0.177 | 0.157 | 404 |
| Rat | Val2 | Pearson P-Val | 1.77E-08 | 4.16E-06 | 4.82E-05 | 8.07E-05 | 1.52E-03 | |
| Rat | Val2 | Spearman | 0.205 | 0.206 | 0.189 | 0.191 | 0.167 | 404 |
| Rat | Val2 | Spearman P-Val | 8.02E-07 | 7.03E-07 | 1.29E-05 | 8.51E-06 | 3.63E-04 | |
| Rat | Val2 | MSE | 1.88 | 1.4 | 1.39 | 1.42 | 1.58 | |
| Rat | Val3 | Pearson | 0.308 | 0.305 | 0.309 | 0.292 | 0.296 | 669 |
| Rat | Val3 | Pearson P-Val | 1.75E-28 | 5.82E-28 | 1.26E-28 | 1.73E-25 | 3.06E-26 | |
| Rat | Val3 | Spearman | 0.301 | 0.293 | 0.295 | 0.282 | 0.282 | 669 |
| Rat | Val3 | Spearman P-Val | 4.84E-27 | 1.55E-25 | 5.44E-26 | 1.24E-23 | 1.26E-23 | |
| Rat | Val3 | MSE | 1.89 | 1.58 | 1.48 | 1.51 | 1.62 | |
| Rat | Test2 | Pearson | 0.259 | 0.219 | 0.224 | 0.204 | 0.232 | 723 |
| Rat | Test2 | Pearson P-Val | 3.20E-21 | 7.05E-15 | 1.19E-15 | 8.84E-13 | 7.59E-17 | |
| Rat | Test2 | Spearman | 0.25 | 0.223 | 0.232 | 0.214 | 0.248 | 723 |
| Rat | Test2 | Spearman P-Val | 8.41E-20 | 1.76E-15 | 8.71E-17 | 4.22E-14 | 1.78E-19 | |
| Rat | Test2 | MSE | 1.92 | 1.5 | 1.47 | 1.5 | 1.63 | |
| Rat | Test3 | Pearson | 0.317 | 0.308 | 0.323 | 0.305 | 0.3 | 1025 |
| Rat | Test3 | Pearson P-Val | 6.76E-47 | 4.06E-44 | 1.48E-48 | 4.64E-43 | 1.57E-41 | |
| Rat | Test3 | Spearman | 0.309 | 0.306 | 0.316 | 0.306 | 0.299 | 1025 |
| Rat | Test3 | Spearman P-Val | 3.72E-44 | 2.41E-43 | 1.40E-46 | 1.63E-43 | 2.05E-41 | |
| Rat | Test3 | MSE | 1.8 | 1.54 | 1.43 | 1.47 | 1.57 | |
| Cow+Pig | Train | Pearson | 0.378 | 0.37 | 0.381 | 0.366 | 0.359 | 360 |
| Cow+Pig | Train | Pearson P-Val | 1.55E-23 | 1.69E-22 | 4.80E-24 | 5.74E-22 | 5.63E-21 | |
| Cow+Pig | Train | Spearman | 0.358 | 0.366 | 0.367 | 0.35 | 0.352 | 360 |
| Cow+Pig | Train | Spearman P-Val | 7.43E-21 | 5.67E-22 | 4.78E-22 | 8.19E-20 | 3.69E-20 | |
| Cow+Pig | Train | MSE | 2.31 | 2.03 | 1.87 | 1.86 | 2 | |

| | | | | | | | | |
|---|---|---|---|---|---|---|---|---|
| Cow+Pig | Validation | Pearson | 0.3 | 0.305 | 0.318 | 0.324 | 0.325 | 73 |
| Cow+Pig | Validation | Pearson P-Val | 4.77E-02 | 3.66E-02 | 1.79E-02 | 1.32E-02 | 1.27E-02 | |
| Cow+Pig | Validation | Spearman | 0.276 | 0.289 | 0.267 | 0.299 | 0.288 | 73 |
| Cow+Pig | Validation | Spearman P-Val | 1.51E-01 | 8.11E-02 | 2.28E-01 | 4.90E-02 | 8.29E-02 | |
| Cow+Pig | Validation | MSE | 1.93 | 1.67 | 1.55 | 1.59 | 1.63 | |
| Cow+Pig | Test3 | Pearson | 0.435 | 0.383 | 0.439 | 0.346 | 0.402 | 77 |
| Cow+Pig | Test3 | Pearson P-Val | 3.52E-06 | 1.89E-04 | 2.56E-06 | 2.24E-03 | 4.63E-05 | |
| Cow+Pig | Test3 | Spearman | 0.456 | 0.427 | 0.481 | 0.376 | 0.451 | 77 |
| Cow+Pig | Test3 | Spearman P-Val | 5.75E-07 | 6.79E-06 | 5.68E-08 | 3.16E-04 | 8.75E-07 | |
| Cow+Pig | Test3 | MSE | 2.19 | 1.9 | 1.76 | 1.76 | 1.92 | |

P-values were adjusted using Bonferroni Correction. MSE: Mean squared error.

**Table S6.** Mean Negative Set Predictions of the 5 best mouse-only trained models

| Species | Group | Best Mouse-Only | Alternate 1 | Alternate 2 | Alternate 3 | Alternate 4 | Number of Peaks |
|---|---|---|---|---|---|---|---|
| Mouse | Train | 0.612 | 0.584 | 0.648 | 0.671 | 0.674 | 18851 |
| Mouse | Validation | 0.619 | 0.6 | 0.664 | 0.682 | 0.684 | 2587 |
| Mouse | Test | 0.609 | 0.59 | 0.654 | 0.673 | 0.675 | 3698 |
| Cow | Train | 0.624 | 0.625 | 0.682 | 0.698 | 0.698 | 17403 |
| Cow | Validation | 0.635 | 0.636 | 0.694 | 0.713 | 0.711 | 2049 |
| Cow | Test | 0.619 | 0.618 | 0.673 | 0.688 | 0.689 | 3720 |
| Cow | Val1 | 0.626 | 0.627 | 0.68 | 0.697 | 0.69 | 539 |
| Cow | Test1 | 0.596 | 0.59 | 0.641 | 0.655 | 0.662 | 1096 |
| Macaque | Train | 0.606 | 0.601 | 0.66 | 0.678 | 0.678 | 16165 |
| Macaque | Validation | 0.61 | 0.611 | 0.666 | 0.687 | 0.688 | 1943 |
| Macaque | Test | 0.604 | 0.599 | 0.662 | 0.676 | 0.678 | 3477 |
| Macaque | Val1 | 0.594 | 0.596 | 0.65 | 0.67 | 0.671 | 542 |
| Macaque | Test1 | 0.588 | 0.582 | 0.648 | 0.659 | 0.664 | 1157 |
| Pig | Train | 0.628 | 0.634 | 0.689 | 0.706 | 0.704 | 14436 |
| Pig | Validation | 0.638 | 0.638 | 0.696 | 0.711 | 0.71 | 1770 |
| Pig | Test | 0.625 | 0.627 | 0.684 | 0.7 | 0.701 | 3057 |
| Pig | Val1 | 0.615 | 0.606 | 0.654 | 0.67 | 0.672 | 453 |
| Pig | Test1 | 0.598 | 0.589 | 0.652 | 0.664 | 0.671 | 940 |
| Rat | Train | 0.656 | 0.665 | 0.721 | 0.738 | 0.732 | 19408 |
| Rat | Validation | 0.676 | 0.691 | 0.748 | 0.764 | 0.755 | 2850 |
| Rat | Test | 0.66 | 0.672 | 0.728 | 0.745 | 0.738 | 4405 |
| Rat | Val1 | 0.708 | 0.754 | 0.812 | 0.814 | 0.8 | 688 |
| Rat | Test1 | 0.698 | 0.748 | 0.802 | 0.813 | 0.797 | 1241 |
| Cow+Pig | Train | 0.658 | 0.684 | 0.724 | 0.735 | 0.737 | 238 |
| Cow+Pig | Validation | 0.682 | 0.737 | 0.771 | 0.807 | 0.766 | 32 |
| Cow+Pig | Test | 0.677 | 0.715 | 0.745 | 0.757 | 0.748 | 86 |

**Table S7.** Performance in Predicting the Direction of Accessibility Differences of the 5 best mouse-only trained models

| Species | Group | Metric | Best Mouse-Only | Alternate 1 | Alternate 2 | Alternate 3 | Alternate 4 | Number of Peaks |
|---|---|---|---|---|---|---|---|---|
| Cow | Val2 | Same Sign Count | 200 | 184 | 184 | 183 | 181 | 330 |
| Cow | Val2 | Same Sign (frac) | 0.606 | 0.558 | 0.558 | 0.555 | 0.548 | |
| Cow | Val2 | Pearson | 0.259 | 0.286 | 0.273 | 0.25 | 0.273 | |
| Cow | Val2 | Pearson P-Val | 3.80E-04 | 2.63E-05 | 9.35E-05 | 8.60E-04 | 9.73E-05 | |
| Cow | Val2 | Spearman | 0.263 | 0.255 | 0.254 | 0.218 | 0.231 | |
| Cow | Val2 | Spearman P-Val | 2.45E-04 | 5.60E-04 | 5.67E-04 | 1.29E-02 | 4.66E-03 | |
| Cow | Val2 | MSE | 0.399 | 0.486 | 0.486 | 0.489 | 0.452 | |
| Cow | Test2 | Same Sign Count | 358 | 327 | 331 | 347 | 333 | 618 |
| Cow | Test2 | Same Sign (frac) | 0.579 | 0.529 | 0.536 | 0.561 | 0.539 | |
| Cow | Test2 | Pearson | 0.201 | 0.18 | 0.228 | 0.195 | 0.187 | |
| Cow | Test2 | Pearson P-Val | 8.80E-05 | 1.33E-03 | 1.95E-06 | 1.99E-04 | 5.89E-04 | |
| Cow | Test2 | Spearman | 0.228 | 0.174 | 0.234 | 0.21 | 0.185 | |
| Cow | Test2 | Spearman P-Val | 1.85E-06 | 2.58E-03 | 8.44E-07 | 2.67E-05 | 7.20E-04 | |
| Cow | Test2 | MSE | 0.389 | 0.463 | 0.433 | 0.442 | 0.424 | |
| Macaque | Val2 | Same Sign Count | 308 | 292 | 295 | 295 | 308 | 520 |
| Macaque | Val2 | Same Sign (frac) | 0.592 | 0.562 | 0.567 | 0.567 | 0.592 | |
| Macaque | Val2 | Pearson | 0.269 | 0.229 | 0.196 | 0.201 | 0.2 | |
| Macaque | Val2 | Pearson P-Val | 9.60E-08 | 2.60E-05 | 1.42E-03 | 7.66E-04 | 8.69E-04 | |
| Macaque | Val2 | Spearman | 0.246 | 0.214 | 0.195 | 0.19 | 0.198 | |
| Macaque | Val2 | Spearman P-Val | 2.51E-06 | 1.66E-04 | 1.57E-03 | 2.48E-03 | 1.07E-03 | |
| Macaque | Val2 | MSE | 0.28 | 0.379 | 0.371 | 0.362 | 0.343 | |
| Macaque | Test2 | Same Sign Count | 466 | 447 | 461 | 449 | 448 | 798 |
| Macaque | Test2 | Same Sign (frac) | 0.584 | 0.56 | 0.578 | 0.563 | 0.561 | |
| Macaque | Test2 | Pearson | 0.212 | 0.195 | 0.202 | 0.167 | 0.121 | |
| Macaque | Test2 | Pearson P-Val | 2.74E-07 | 5.95E-06 | 1.59E-06 | 4.28E-04 | 1.19E-01 | |
| Macaque | Test2 | Spearman | 0.202 | 0.165 | 0.183 | 0.151 | 0.102 | |

| | | | | | | | | |
|---|---|---|---|---|---|---|---|---|
| Macaque | Test2 | Spearman P-Val | 1.64E-06 | 5.43E-04 | 3.87E-05 | 3.89E-03 | 7.55E-01 | |
| Macaque | Test2 | MSE | 0.294 | 0.375 | 0.355 | 0.374 | 0.369 | |
| Pig | Val2 | Same Sign Count | 146 | 134 | 131 | 129 | 132 | 276 |
| Pig | Val2 | Same Sign (frac) | 0.529 | 0.486 | 0.475 | 0.467 | 0.478 | |
| Pig | Val2 | Pearson | 0.109 | 0.0665 | 0.0635 | 0.0585 | 0.0225 | |
| Pig | Val2 | Pearson P-Val | 1.39E+01 | 5.41E+01 | 5.87E+01 | 6.65E+01 | 1.42E+02 | |
| Pig | Val2 | Spearman | 0.133 | 0.0678 | 0.0604 | 0.0572 | 0.0146 | |
| Pig | Val2 | Spearman P-Val | 5.45E+00 | 5.23E+01 | 6.35E+01 | 6.88E+01 | 1.62E+02 | |
| Pig | Val2 | MSE | 0.358 | 0.503 | 0.5 | 0.492 | 0.471 | |
| Pig | Test2 | Same Sign Count | 285 | 264 | 271 | 277 | 275 | 544 |
| Pig | Test2 | Same Sign (frac) | 0.524 | 0.485 | 0.498 | 0.509 | 0.506 | |
| Pig | Test2 | Pearson | 0.249 | 0.205 | 0.224 | 0.188 | 0.196 | |
| Pig | Test2 | Pearson P-Val | 7.55E-07 | 2.79E-04 | 2.48E-05 | 1.99E-03 | 8.02E-04 | |
| Pig | Test2 | Spearman | 0.195 | 0.137 | 0.151 | 0.148 | 0.138 | |
| Pig | Test2 | Spearman P-Val | 8.94E-04 | 2.63E-01 | 8.33E-02 | 1.06E-01 | 2.48E-01 | |
| Pig | Test2 | MSE | 0.343 | 0.46 | 0.431 | 0.429 | 0.402 | |
| Rat | Val2 | Same Sign Count | 483 | 471 | 474 | 457 | 453 | 796 |
| Rat | Val2 | Same Sign (frac) | 0.607 | 0.592 | 0.595 | 0.574 | 0.569 | |
| Rat | Val2 | Pearson | 0.159 | 0.172 | 0.169 | 0.162 | 0.153 | |
| Rat | Val2 | Pearson P-Val | 1.33E-03 | 2.15E-04 | 3.11E-04 | 9.06E-04 | 2.81E-03 | |
| Rat | Val2 | Spearman | 0.204 | 0.194 | 0.195 | 0.166 | 0.172 | |
| Rat | Val2 | Spearman P-Val | 1.34E-06 | 6.53E-06 | 5.94E-06 | 4.81E-04 | 2.18E-04 | |
| Rat | Val2 | MSE | 0.205 | 0.25 | 0.24 | 0.244 | 0.224 | |
| Rat | Test2 | Same Sign Count | 789 | 812 | 794 | 785 | 779 | 1440 |
| Rat | Test2 | Same Sign (frac) | 0.549 | 0.565 | 0.552 | 0.546 | 0.542 | |
| Rat | Test2 | Pearson | 0.177 | 0.155 | 0.151 | 0.135 | 0.13 | |
| Rat | Test2 | Pearson P-Val | 2.53E-09 | 7.42E-07 | 1.62E-06 | 5.33E-05 | 1.57E-04 | |
| Rat | Test2 | Spearman | 0.153 | 0.145 | 0.138 | 0.124 | 0.119 | |
| Rat | Test2 | Spearman P-Val | 1.17E-06 | 6.92E-06 | 2.76E-05 | 5.05E-04 | 1.23E-03 | |

| | | | | | | | |
|---|---|---|---|---|---|---|---|
| Rat | Test2 | MSE | 0.201 | 0.25 | 0.239 | 0.25 | 0.232 |

P-values were adjusted using Bonferroni Correction. MSE: Mean squared error.

**Table S8.** Performance of the best log-transformed vs. quantile normalized (QN) vs. extended quantile normalized (EQN) vs. 2000 bp models

| Species | Group | Metric | Best Log | QN | EQN | 2000 bp | Number of Peaks QN / Others |
|---|---|---|---|---|---|---|---|
| Mouse | Test | Pearson | 0.496 | 0.466 | 0.479 | 0.464 | 3485 / 3776 |
| Mouse | Test | Pearson P-Val | 0.00E+00 | 0.00E+00 | 0.00E+00 | 0.00E+00 | |
| Mouse | Test | Spearman | 0.502 | 0.473 | 0.477 | 0.464 | 3485 / 3776 |
| Mouse | Test | Spearman P-Val | 0.00E+00 | 0.00E+00 | 0.00E+00 | 0.00E+00 | |
| Mouse | Test | MSE | 1.44 | 1.2 | 1.05 | 1.52 | |
| Cow | Test | Pearson | 0.385 | 0.319 | 0.355 | 0.422 | 1124 / 1284 |
| Cow | Test | Pearson P-Val | 2.67E-89 | 6.43E-52 | 1.37E-74 | 1.88E-109 | |
| Cow | Test | Spearman | 0.391 | 0.322 | 0.366 | 0.43 | 1124 / 1284 |
| Cow | Test | Spearman P-Val | 1.77E-92 | 5.32E-53 | 9.32E-80 | 2.47E-114 | |
| Cow | Test | MSE | 2.32 | 1.2 | 1.14 | 2.52 | |
| Cow | Test2 | Pearson | 0.364 | 0.303 | 0.36 | 0.37 | 302 / 337 |
| Cow | Test2 | Pearson P-Val | 3.37E-20 | 5.50E-12 | 1.33E-19 | 2.35E-21 | |
| Cow | Test2 | Spearman | 0.364 | 0.316 | 0.369 | 0.38 | 302 / 337 |
| Cow | Test2 | Spearman P-Val | 3.46E-20 | 3.82E-13 | 1.17E-20 | 1.43E-22 | |
| Cow | Test2 | MSE | 2.28 | 1.01 | 0.944 | 2.61 | |
| Cow | Test3 | Pearson | 0.319 | 0.264 | 0.283 | 0.371 | 597 / 694 |
| Cow | Test3 | Pearson P-Val | 8.43E-32 | 3.84E-18 | 1.96E-24 | 1.60E-44 | |
| Cow | Test3 | Spearman | 0.321 | 0.263 | 0.284 | 0.37 | 597 / 694 |
| Cow | Test3 | Spearman P-Val | 2.20E-32 | 5.02E-18 | 1.06E-24 | 2.44E-44 | |
| Cow | Test3 | MSE | 2.22 | 1.22 | 1.2 | 2.32 | |
| Macaque | Test | Pearson | 0.327 | 0.231 | 0.371 | 0.348 | 1681 / 2624 |
| Macaque | Test | Pearson P-Val | 2.07E-128 | 1.69E-39 | 1.00E-168 | 1.73E-147 | |
| Macaque | Test | Spearman | 0.346 | 0.225 | 0.372 | 0.359 | 1681 / 2624 |
| Macaque | Test | Spearman P-Val | 2.89E-145 | 1.52E-37 | 4.00E-169 | 9.80E-158 | |
| Macaque | Test | MSE | 1.59 | 1.18 | 1.25 | 1.66 | |
| Macaque | Test2 | Pearson | 0.289 | 0.216 | 0.325 | 0.289 | 354 / 472 |
| Macaque | Test2 | Pearson P-Val | 2.23E-17 | 1.28E-06 | 4.00E-22 | 1.36E-17 | |
| Macaque | Test2 | Spearman | 0.274 | 0.219 | 0.305 | 0.28 | 354 / 472 |
| Macaque | Test2 | Spearman P-Val | 2.24E-15 | 7.77E-07 | 2.59E-19 | 1.97E-16 | |
| Macaque | Test2 | MSE | 1.58 | 1.04 | 1.06 | 1.73 | |
| Macaque | Test3 | Pearson | 0.342 | 0.226 | 0.417 | 0.362 | 777 / 1263 |

| | | | | | | | |
|---|---|---|---|---|---|---|---|
| Macaque | Test3 | Pearson P-Val | 6.17E-68 | 3.89E-17 | 2.57E-104 | 3.57E-77 | |
| Macaque | Test3 | Spearman | 0.369 | 0.219 | 0.424 | 0.38 | 777 / 1263 |
| Macaque | Test3 | Spearman P-Val | 5.09E-80 | 5.16E-16 | 3.31E-108 | 1.74E-85 | |
| Macaque | Test3 | MSE | 1.67 | 1.27 | 1.37 | 1.69 | |
| Pig | Test | Pearson | 0.388 | 0.354 | 0.349 | 0.427 | 1627 / 1627 |
| Pig | Test | Pearson P-Val | 2.24E-115 | 2.71E-94 | 4.26E-91 | 4.36E-142 | |
| Pig | Test | Spearman | 0.394 | 0.358 | 0.358 | 0.43 | 1627 / 1627 |
| Pig | Test | Spearman P-Val | 2.51E-119 | 9.91E-97 | 2.31E-96 | 1.86E-144 | |
| Pig | Test | MSE | 2.25 | 1.51 | 1.51 | 2.39 | |
| Pig | Test2 | Pearson | 0.351 | 0.275 | 0.293 | 0.385 | 293 / 293 |
| Pig | Test2 | Pearson P-Val | 4.29E-16 | 2.30E-09 | 1.41E-10 | 3.51E-20 | |
| Pig | Test2 | Spearman | 0.342 | 0.259 | 0.28 | 0.371 | 293 / 293 |
| Pig | Test2 | Spearman P-Val | 3.58E-15 | 3.81E-08 | 1.51E-09 | 1.47E-18 | |
| Pig | Test2 | MSE | 2.42 | 1.49 | 1.41 | 2.66 | |
| Pig | Test3 | Pearson | 0.337 | 0.339 | 0.329 | 0.376 | 716 / 716 |
| Pig | Test3 | Pearson P-Val | 4.27E-37 | 1.62E-37 | 4.39E-35 | 2.78E-47 | |
| Pig | Test3 | Spearman | 0.337 | 0.328 | 0.329 | 0.381 | 716 / 716 |
| Pig | Test3 | Spearman P-Val | 3.89E-37 | 5.03E-35 | 4.69E-35 | 1.13E-48 | |
| Pig | Test3 | MSE | 2.16 | 1.55 | 1.59 | 2.22 | |
| Rat | Test | Pearson | 0.333 | 0.342 | 0.341 | 0.337 | 2462 / 2576 |
| Rat | Test | Pearson P-Val | 5.67E-131 | 6.60E-133 | 4.87E-138 | 8.60E-135 | |
| Rat | Test | Spearman | 0.332 | 0.336 | 0.335 | 0.331 | 2462 / 2576 |
| Rat | Test | Spearman P-Val | 1.07E-130 | 3.71E-128 | 3.82E-133 | 5.13E-130 | |
| Rat | Test | MSE | 1.82 | 1.19 | 1.06 | 1.94 | |
| Rat | Test2 | Pearson | 0.259 | 0.273 | 0.267 | 0.274 | 705 / 723 |
| Rat | Test2 | Pearson P-Val | 3.20E-21 | 3.41E-23 | 1.65E-22 | 3.03E-24 | |
| Rat | Test2 | Spearman | 0.25 | 0.261 | 0.256 | 0.259 | 705 / 723 |
| Rat | Test2 | Spearman P-Val | 8.41E-20 | 4.44E-21 | 1.50E-20 | 1.44E-21 | |
| Rat | Test2 | MSE | 1.92 | 1.06 | 0.992 | 2.12 | |
| Rat | Test3 | Pearson | 0.317 | 0.334 | 0.324 | 0.324 | 951 / 1025 |
| Rat | Test3 | Pearson P-Val | 6.76E-47 | 1.34E-48 | 2.84E-48 | 9.79E-49 | |
| Rat | Test3 | Spearman | 0.309 | 0.322 | 0.31 | 0.315 | 951 / 1025 |
| Rat | Test3 | Spearman P-Val | 3.72E-44 | 8.39E-45 | 6.30E-44 | 9.01E-46 | |
| Rat | Test3 | MSE | 1.8 | 1.26 | 1.12 | 1.87 | |
| Cow+Pig | Test3 | Pearson | 0.435 | 0.351 | 0.422 | 0.459 | 66 / 77 |

| | | | | | | | |
|---|---|---|---|---|---|---|---|
| Cow+Pig | Test3 | Pearson P-Val | 3.52E-06 | 7.20E-03 | 1.49E-05 | 2.15E-07 | |
| Cow+Pig | Test3 | Spearman | 0.456 | 0.425 | 0.455 | 0.484 | 66 / 77 |
| Cow+Pig | Test3 | Spearman P-Val | 5.75E-07 | 7.61E-05 | 9.48E-07 | 2.12E-08 | |
| Cow+Pig | Test3 | MSE | 2.19 | 1.2 | 1.19 | 2.31 | |

P-values were adjusted using Bonferroni Correction. Note Best Log and Best Mouse-Only are the same model. MSE: Mean squared error.

**Table S9.** Performance in Predicting the Direction of Accessibility Difference of the best mouse-only log-transformed vs. quantile normalized (QN) vs. extended quantile normalized (EQN) vs. 2000 bp models

| Species | Metric | Best Log | QN | EQN | 2000 bp | Number of Peaks QN / Others |
|---|---|---|---|---|---|---|
| Cow | Same Sign Count | 358 | 332 | 380 | 327 | 584 / 618 |
| Cow | Same Sign (frac) | 0.579 | 0.568 | 0.615 | 0.529 | |
| Cow | Pearson | 0.201 | 0.302 | 0.202 | 0.199 | |
| Cow | Pearson P-Val | 8.80E-05 | 1.72E-11 | 1.26E-04 | 5.93E-05 | |
| Cow | Spearman | 0.228 | 0.277 | 0.225 | 0.226 | |
| Cow | Spearman P-Val | 1.85E-06 | 1.81E-09 | 4.83E-06 | 1.38E-06 | |
| Cow | MSE | 0.389 | 0.722 | 0.53 | 0.382 | |
| Macaque | Same Sign Count | 466 | 397 | 446 | 461 | 670 / 798 |
| Macaque | Same Sign (frac) | 0.584 | 0.593 | 0.559 | 0.578 | |
| Macaque | Pearson | 0.212 | 0.22 | 0.161 | 0.186 | |
| Macaque | Pearson P-Val | 2.74E-07 | 1.62E-06 | 1.54E-03 | 1.20E-05 | |
| Macaque | Spearman | 0.202 | 0.214 | 0.141 | 0.181 | |
| Macaque | Spearman P-Val | 1.64E-06 | 4.31E-06 | 1.98E-02 | 2.58E-05 | |
| Macaque | MSE | 0.294 | 0.867 | 0.557 | 0.289 | |
| Pig | Same Sign Count | 285 | 291 | 274 | 282 | 544 / 544 |
| Pig | Same Sign (frac) | 0.524 | 0.535 | 0.504 | 0.518 | |
| Pig | Pearson | 0.249 | 0.201 | 0.183 | 0.239 | |
| Pig | Pearson P-Val | 7.55E-07 | 4.46E-04 | 5.28E-03 | 1.74E-06 | |
| Pig | Spearman | 0.195 | 0.144 | 0.131 | 0.173 | |
| Pig | Spearman P-Val | 8.94E-04 | 1.55E-01 | 6.40E-01 | 4.77E-03 | |
| Pig | MSE | 0.343 | 0.688 | 0.594 | 0.323 | |
| Rat | Same Sign Count | 789 | 820 | 791 | 813 | 1406 / 1440 |
| Rat | Same Sign (frac) | 0.549 | 0.583 | 0.55 | 0.565 | |
| Rat | Pearson | 0.177 | 0.282 | 0.173 | 0.182 | |
| Rat | Pearson P-Val | 2.53E-09 | 7.09E-25 | 1.30E-08 | 3.96E-10 | |
| Rat | Spearman | 0.153 | 0.265 | 0.152 | 0.165 | |
| Rat | Spearman P-Val | 1.17E-06 | 9.21E-22 | 2.19E-06 | 3.26E-08 | |
| Rat | MSE | 0.201 | 0.851 | 0.434 | 0.191 | |

P-values were adjusted using Bonferroni Correction. Note Best Log and Best Mouse-Only are the same model. MSE: Mean squared error.

**Table S10.** Mean Negative Set Predictions of the best mouse-only log-transformed vs. quantile normalized (QN) vs. extended quantile normalized (EQN) vs. 2000 bp models

| Species | Group | Best Log | QN | EQN | 2000 bp | Number of Peaks |
|---|---|---|---|---|---|---|
| Mouse | Test | 0.609 | 0.495 | 0.541 | 0.645 | 3698 |
| Cow | Test | 0.619 | 0.512 | 0.568 | 0.658 | 3720 |
| Cow | Test1 | 0.596 | 0.454 | 0.52 | 0.642 | 1096 |
| Macaque | Test | 0.604 | 0.473 | 0.542 | 0.648 | 3477 |
| Macaque | Test1 | 0.588 | 0.444 | 0.515 | 0.638 | 1157 |
| Pig | Test | 0.625 | 0.527 | 0.594 | 0.658 | 3057 |
| Pig | Test1 | 0.598 | 0.469 | 0.538 | 0.638 | 940 |
| Rat | Test | 0.66 | 0.623 | 0.676 | 0.675 | 4405 |
| Rat | Test1 | 0.698 | 0.758 | 0.804 | 0.71 | 1241 |
| Cow+Pig | Test1 | 0.677 | 0.623 | 0.717 | 0.708 | 86 |

**Table S11.** Mean Negative Set Predictions of the best mouse-only vs. 3-species vs. 5-species models

| Species | Group | Mouse-Only | 3-Species | 5-Species | Number of Peaks |
|---|---|---|---|---|---|
| Mouse | Train | 0.612 | 0.403 | 0.401 | 18851 |
| Mouse | Validation | 0.619 | 0.46 | 0.429 | 2587 |
| Mouse | Test | 0.609 | 0.446 | 0.424 | 3698 |
| Cow | Train | 0.624 | 0.498 | 0.422 | 17403 |
| Cow | Validation | 0.635 | 0.52 | 0.48 | 2049 |
| Cow | Test | 0.619 | 0.482 | 0.46 | 3720 |
| Cow | Val1 | 0.626 | 0.472 | 0.427 | 539 |
| Cow | Test1 | 0.596 | 0.416 | 0.393 | 1096 |
| Macaque | Train | 0.606 | 0.373 | 0.379 | 16165 |
| Macaque | Validation | 0.61 | 0.428 | 0.412 | 1943 |
| Macaque | Test | 0.604 | 0.432 | 0.423 | 3477 |
| Macaque | Val1 | 0.594 | 0.387 | 0.372 | 542 |
| Macaque | Test1 | 0.588 | 0.386 | 0.379 | 1157 |
| Pig | Train | 0.628 | 0.493 | 0.393 | 14436 |
| Pig | Validation | 0.638 | 0.499 | 0.447 | 1770 |
| Pig | Test | 0.625 | 0.489 | 0.451 | 3057 |
| Pig | Val1 | 0.615 | 0.422 | 0.387 | 453 |
| Pig | Test1 | 0.598 | 0.401 | 0.374 | 940 |
| Rat | Train | 0.656 | 0.468 | 0.455 | 19408 |
| Rat | Validation | 0.676 | 0.59 | 0.537 | 2850 |
| Rat | Test | 0.66 | 0.555 | 0.516 | 4405 |
| Rat | Val1 | 0.708 | 0.722 | 0.655 | 688 |
| Rat | Test1 | 0.698 | 0.68 | 0.634 | 1241 |
| Cow+Pig | Train | 0.658 | 0.596 | 0.472 | 238 |
| Cow+Pig | Validation | 0.682 | 0.685 | 0.576 | 32 |
| Cow+Pig | Test | 0.677 | 0.626 | 0.59 | 86 |

**Table S12.** Performance of the best mouse-only vs. 3-species vs. 5-species models Performance in Predicting the Direction of Accessibility Difference of

| Species | Group | Metric | Mouse-Only | 3-Species | 5-Species | Number of Peaks |
|---|---|---|---|---|---|---|
| Mouse | Train | Pearson | 0.494 | 0.644 | 0.645 | 16250 |
| Mouse | Train | Pearson P-Val | 0.00E+00 | 0.00E+00 | 0.00E+00 | |
| Mouse | Train | Spearman | 0.498 | 0.64 | 0.639 | 16250 |
| Mouse | Train | Spearman P-Val | 0.00E+00 | 0.00E+00 | 0.00E+00 | |
| Mouse | Train | MSE | 1.39 | 0.521 | 0.622 | |
| Mouse | Validation | Pearson | 0.483 | 0.584 | 0.597 | 2016 |
| Mouse | Validation | Pearson P-Val | 3.49E-232 | 0.00E+00 | 0.00E+00 | |
| Mouse | Validation | Spearman | 0.491 | 0.582 | 0.588 | 2016 |
| Mouse | Validation | Spearman P-Val | 8.59E-242 | 0.00E+00 | 0.00E+00 | |
| Mouse | Validation | MSE | 1.42 | 0.638 | 0.699 | |
| Mouse | Test | Pearson | 0.496 | 0.6 | 0.628 | 3776 |
| Mouse | Test | Pearson P-Val | 0.00E+00 | 0.00E+00 | 0.00E+00 | |
| Mouse | Test | Spearman | 0.502 | 0.601 | 0.626 | 3776 |
| Mouse | Test | Spearman P-Val | 0.00E+00 | 0.00E+00 | 0.00E+00 | |
| Mouse | Test | MSE | 1.44 | 0.67 | 0.737 | |
| Cow | Train | Pearson | 0.406 | 0.429 | 0.518 | 7687 |
| Cow | Train | Pearson P-Val | 0.00E+00 | 0.00E+00 | 0.00E+00 | |
| Cow | Train | Spearman | 0.401 | 0.429 | 0.518 | 7687 |
| Cow | Train | Spearman P-Val | 0.00E+00 | 0.00E+00 | 0.00E+00 | |
| Cow | Train | MSE | 2.27 | 1.11 | 1.05 | |
| Cow | Validation | Pearson | 0.4 | 0.439 | 0.506 | 1162 |
| Cow | Validation | Pearson P-Val | 3.61E-88 | 3.41E-108 | 3.98E-149 | |
| Cow | Validation | Spearman | 0.39 | 0.443 | 0.504 | 1162 |
| Cow | Validation | Spearman P-Val | 3.06E-83 | 3.21E-110 | 2.64E-148 | |
| Cow | Validation | MSE | 2.15 | 1.09 | 1.18 | |
| Cow | Test | Pearson | 0.385 | 0.42 | 0.475 | 1284 |
| Cow | Test | Pearson P-Val | 1.34E-89 | 1.58E-108 | 2.35E-142 | |
| Cow | Test | Spearman | 0.391 | 0.432 | 0.485 | 1284 |
| Cow | Test | Spearman P-Val | 8.86E-93 | 4.19E-115 | 9.78E-150 | |
| Cow | Test | MSE | 2.32 | 1.14 | 1.26 | |
| Cow | Val2 | Pearson | 0.547 | 0.546 | 0.618 | 194 |
| Cow | Val2 | Pearson P-Val | 1.23E-29 | 1.81E-29 | 2.70E-40 | |

| | | | | | | |
|---|---|---|---|---|---|---|
| Cow | Val2 | Spearman | 0.54 | 0.542 | 0.59 | 194 |
| Cow | Val2 | Spearman P-Val | 1.09E-28 | 5.15E-29 | 7.98E-36 | |
| Cow | Val2 | MSE | 2.26 | 0.911 | 1.04 | |
| Cow | Val3 | Pearson | 0.256 | 0.362 | 0.423 | 556 |
| Cow | Val3 | Pearson P-Val | 4.12E-16 | 9.05E-34 | 1.35E-47 | |
| Cow | Val3 | Spearman | 0.263 | 0.363 | 0.418 | 556 |
| Cow | Val3 | Spearman P-Val | 4.08E-17 | 6.92E-34 | 2.40E-46 | |
| Cow | Val3 | MSE | 2.12 | 1.21 | 1.28 | |
| Cow | Test2 | Pearson | 0.364 | 0.416 | 0.446 | 337 |
| Cow | Test2 | Pearson P-Val | 1.69E-20 | 1.39E-27 | 2.79E-32 | |
| Cow | Test2 | Spearman | 0.364 | 0.44 | 0.47 | 337 |
| Cow | Test2 | Spearman P-Val | 1.73E-20 | 2.68E-31 | 2.21E-36 | |
| Cow | Test2 | MSE | 2.28 | 0.936 | 1.09 | |
| Cow | Test3 | Pearson | 0.319 | 0.382 | 0.424 | 694 |
| Cow | Test3 | Pearson P-Val | 4.22E-32 | 2.27E-47 | 1.02E-59 | |
| Cow | Test3 | Spearman | 0.321 | 0.379 | 0.417 | 694 |
| Cow | Test3 | Spearman P-Val | 1.10E-32 | 1.19E-46 | 2.33E-57 | |
| Cow | Test3 | MSE | 2.22 | 1.14 | 1.27 | |
| Macaque | Train | Pearson | 0.361 | 0.558 | 0.566 | 12066 |
| Macaque | Train | Pearson P-Val | 0.00E+00 | 0.00E+00 | 0.00E+00 | |
| Macaque | Train | Spearman | 0.374 | 0.556 | 0.559 | 12066 |
| Macaque | Train | Spearman P-Val | 0.00E+00 | 0.00E+00 | 0.00E+00 | |
| Macaque | Train | MSE | 1.6 | 0.62 | 0.721 | |
| Macaque | Validation | Pearson | 0.355 | 0.488 | 0.498 | 1788 |
| Macaque | Validation | Pearson P-Val | 3.57E-104 | 4.25E-211 | 2.90E-221 | |
| Macaque | Validation | Spearman | 0.366 | 0.481 | 0.485 | 1788 |
| Macaque | Validation | Spearman P-Val | 1.46E-111 | 1.49E-204 | 9.40E-209 | |
| Macaque | Validation | MSE | 1.56 | 0.723 | 0.823 | |
| Macaque | Test | Pearson | 0.327 | 0.488 | 0.519 | 2624 |
| Macaque | Test | Pearson P-Val | 2.07E-128 | 0.00E-02 | 0.00E+00 | |
| Macaque | Test | Spearman | 0.346 | 0.482 | 0.512 | 2624 |
| Macaque | Test | Spearman P-Val | 2.89E-145 | 3.30E-301 | 0.00E+00 | |
| Macaque | Test | MSE | 1.59 | 0.788 | 0.855 | |
| Macaque | Val2 | Pearson | 0.437 | 0.531 | 0.561 | 331 |
| Macaque | Val2 | Pearson P-Val | 5.34E-30 | 1.90E-47 | 3.59E-54 | |

| | | | | | | |
|---|---|---|---|---|---|---|
| Macaque | Val2 | Spearman | 0.431 | 0.484 | 0.512 | 331 |
| Macaque | Val2 | Spearman P-Val | 4.72E-29 | 3.67E-38 | 1.74E-43 | |
| Macaque | Val2 | MSE | 1.55 | 0.588 | 0.652 | |
| Macaque | Val3 | Pearson | 0.33 | 0.461 | 0.466 | 887 |
| Macaque | Val3 | Pearson P-Val | 5.18E-44 | 5.08E-92 | 2.46E-94 | |
| Macaque | Val3 | Spearman | 0.345 | 0.456 | 0.461 | 887 |
| Macaque | Val3 | Spearman P-Val | 1.49E-48 | 5.00E-90 | 7.30E-92 | |
| Macaque | Val3 | MSE | 1.64 | 0.835 | 0.938 | |
| Macaque | Test2 | Pearson | 0.289 | 0.444 | 0.441 | 472 |
| Macaque | Test2 | Pearson P-Val | 2.23E-17 | 7.44E-45 | 3.73E-44 | |
| Macaque | Test2 | Spearman | 0.274 | 0.425 | 0.405 | 472 |
| Macaque | Test2 | Spearman P-Val | 2.24E-15 | 8.59E-41 | 1.76E-36 | |
| Macaque | Test2 | MSE | 1.58 | 0.692 | 0.782 | |
| Macaque | Test3 | Pearson | 0.342 | 0.518 | 0.546 | 1263 |
| Macaque | Test3 | Pearson P-Val | 6.17E-68 | 1.87E-171 | 3.85E-194 | |
| Macaque | Test3 | Spearman | 0.369 | 0.515 | 0.544 | 1263 |
| Macaque | Test3 | Spearman P-Val | 5.09E-80 | 6.82E-169 | 1.11E-192 | |
| Macaque | Test3 | MSE | 1.67 | 0.845 | 0.903 | |
| Pig | Train | Pearson | 0.355 | 0.388 | 0.49 | 7762 |
| Pig | Train | Pearson P-Val | 0.00E+00 | 0.00E+00 | 0.00E+00 | |
| Pig | Train | Spearman | 0.347 | 0.384 | 0.489 | 7762 |
| Pig | Train | Spearman P-Val | 0.00E+00 | 0.00E+00 | 0.00E+00 | |
| Pig | Train | MSE | 2.26 | 1.25 | 1.14 | |
| Pig | Validation | Pearson | 0.394 | 0.393 | 0.456 | 992 |
| Pig | Validation | Pearson P-Val | 1.73E-72 | 3.36E-72 | 3.07E-100 | |
| Pig | Validation | Spearman | 0.394 | 0.402 | 0.464 | 992 |
| Pig | Validation | Spearman P-Val | 1.04E-72 | 3.89E-76 | 2.28E-104 | |
| Pig | Validation | MSE | 2.08 | 1.18 | 1.23 | |
| Pig | Test | Pearson | 0.388 | 0.404 | 0.457 | 1627 |
| Pig | Test | Pearson P-Val | 1.12E-115 | 3.20E-126 | 2.63E-165 | |
| Pig | Test | Spearman | 0.394 | 0.406 | 0.459 | 1627 |
| Pig | Test | Spearman P-Val | 1.26E-119 | 1.22E-127 | 3.84E-167 | |
| Pig | Test | MSE | 2.25 | 1.28 | 1.34 | |
| Pig | Val2 | Pearson | 0.456 | 0.42 | 0.54 | 164 |
| Pig | Val2 | Pearson P-Val | 3.03E-16 | 1.89E-13 | 3.07E-24 | |

| | | | | | | |
|---|---|---|---|---|---|---|
| Pig | Val2 | Spearman | 0.45 | 0.414 | 0.511 | 164 |
| Pig | Val2 | Spearman P-Val | 8.97E-16 | 4.90E-13 | 3.52E-21 | |
| Pig | Val2 | MSE | 2.14 | 1.03 | 1.01 | |
| Pig | Val3 | Pearson | 0.274 | 0.338 | 0.403 | 475 |
| Pig | Val3 | Pearson P-Val | 7.54E-16 | 6.81E-25 | 2.57E-36 | |
| Pig | Val3 | Spearman | 0.269 | 0.325 | 0.401 | 475 |
| Pig | Val3 | Spearman P-Val | 2.89E-15 | 9.31E-23 | 4.59E-36 | |
| Pig | Val3 | MSE | 2.08 | 1.32 | 1.37 | |
| Pig | Test2 | Pearson | 0.351 | 0.34 | 0.404 | 293 |
| Pig | Test2 | Pearson P-Val | 2.15E-16 | 2.26E-15 | 1.88E-22 | |
| Pig | Test2 | Spearman | 0.342 | 0.317 | 0.372 | 293 |
| Pig | Test2 | Spearman P-Val | 1.79E-15 | 4.17E-13 | 1.25E-18 | |
| Pig | Test2 | MSE | 2.42 | 1.21 | 1.22 | |
| Pig | Test3 | Pearson | 0.337 | 0.375 | 0.437 | 716 |
| Pig | Test3 | Pearson P-Val | 2.13E-37 | 4.69E-47 | 9.26E-66 | |
| Pig | Test3 | Spearman | 0.337 | 0.37 | 0.428 | 716 |
| Pig | Test3 | Spearman P-Val | 1.94E-37 | 1.08E-45 | 5.22E-63 | |
| Pig | Test3 | MSE | 2.16 | 1.33 | 1.4 | |
| Rat | Train | Pearson | 0.327 | 0.454 | 0.44 | 13365 |
| Rat | Train | Pearson P-Val | 0.00E+00 | 0.00E+00 | 0.00E+00 | |
| Rat | Train | Spearman | 0.331 | 0.461 | 0.446 | 13365 |
| Rat | Train | Spearman P-Val | 0.00E+00 | 0.00E+00 | 0.00E+00 | |
| Rat | Train | MSE | 1.89 | 0.653 | 0.746 | |
| Rat | Validation | Pearson | 0.301 | 0.366 | 0.37 | 1533 |
| Rat | Validation | Pearson P-Val | 4.55E-63 | 1.49E-95 | 2.68E-98 | |
| Rat | Validation | Spearman | 0.304 | 0.374 | 0.378 | 1533 |
| Rat | Validation | Spearman P-Val | 1.10E-64 | 2.38E-100 | 6.37E-103 | |
| Rat | Validation | MSE | 1.84 | 0.758 | 0.825 | |
| Rat | Test | Pearson | 0.333 | 0.419 | 0.424 | 2576 |
| Rat | Test | Pearson P-Val | 2.83E-131 | 6.25E-216 | 1.81E-221 | |
| Rat | Test | Spearman | 0.332 | 0.419 | 0.422 | 2576 |
| Rat | Test | Spearman P-Val | 5.37E-131 | 2.72E-216 | 1.24E-219 | |
| Rat | Test | MSE | 1.82 | 0.813 | 0.843 | |
| Rat | Val2 | Pearson | 0.226 | 0.3 | 0.332 | 404 |
| Rat | Val2 | Pearson P-Val | 8.83E-09 | 2.86E-16 | 3.35E-20 | |

| | | | | | | |
|---|---|---|---|---|---|---|
| Rat | Val2 | Spearman | 0.205 | 0.285 | 0.339 | 404 |
| Rat | Val2 | Spearman P-Val | 4.01E-07 | 1.36E-14 | 3.85E-21 | |
| Rat | Val2 | MSE | 1.88 | 0.615 | 0.667 | |
| Rat | Val3 | Pearson | 0.308 | 0.383 | 0.386 | 669 |
| Rat | Val3 | Pearson P-Val | 8.73E-29 | 4.03E-46 | 1.01E-46 | |
| Rat | Val3 | Spearman | 0.301 | 0.391 | 0.386 | 669 |
| Rat | Val3 | Spearman P-Val | 2.42E-27 | 3.72E-48 | 9.41E-47 | |
| Rat | Val3 | MSE | 1.89 | 0.849 | 0.893 | |
| Rat | Test2 | Pearson | 0.259 | 0.371 | 0.367 | 723 |
| Rat | Test2 | Pearson P-Val | 1.60E-21 | 2.67E-46 | 2.47E-45 | |
| Rat | Test2 | Spearman | 0.25 | 0.375 | 0.361 | 723 |
| Rat | Test2 | Spearman P-Val | 4.20E-20 | 1.94E-47 | 8.06E-44 | |
| Rat | Test2 | MSE | 1.92 | 0.678 | 0.742 | |
| Rat | Test3 | Pearson | 0.317 | 0.393 | 0.403 | 1025 |
| Rat | Test3 | Pearson P-Val | 3.38E-47 | 1.03E-74 | 4.56E-79 | |
| Rat | Test3 | Spearman | 0.309 | 0.384 | 0.398 | 1025 |
| Rat | Test3 | Spearman P-Val | 1.86E-44 | 3.40E-71 | 7.61E-77 | |
| Rat | Test3 | MSE | 1.8 | 0.884 | 0.898 | |
| Cow+Pig | Train | Pearson | 0.378 | 0.451 | 0.53 | 360 |
| Cow+Pig | Train | Pearson P-Val | 7.73E-24 | 1.93E-35 | 2.63E-51 | |
| Cow+Pig | Train | Spearman | 0.358 | 0.428 | 0.521 | 360 |
| Cow+Pig | Train | Spearman P-Val | 3.71E-21 | 2.02E-31 | 2.98E-49 | |
| Cow+Pig | Train | MSE | 2.31 | 1.31 | 1.24 | |
| Cow+Pig | Validation | Pearson | 0.3 | 0.397 | 0.451 | 73 |
| Cow+Pig | Validation | Pearson P-Val | 2.39E-02 | 6.99E-05 | 1.07E-06 | |
| Cow+Pig | Validation | Spearman | 0.276 | 0.341 | 0.371 | 73 |
| Cow+Pig | Validation | Spearman P-Val | 7.53E-02 | 2.60E-03 | 3.99E-04 | |
| Cow+Pig | Validation | MSE | 1.93 | 1.02 | 1.1 | |
| Cow+Pig | Test3 | Pearson | 0.435 | 0.396 | 0.436 | 77 |
| Cow+Pig | Test3 | Pearson P-Val | 1.76E-06 | 3.60E-05 | 1.57E-06 | |
| Cow+Pig | Test3 | Spearman | 0.456 | 0.433 | 0.475 | 77 |
| Cow+Pig | Test3 | Spearman P-Val | 2.88E-07 | 2.02E-06 | 4.70E-08 | |
| Cow+Pig | Test3 | MSE | 2.19 | 1.19 | 1.35 | |

P-values were adjusted using Bonferroni Correction. MSE: Mean squared error.

**Table S13.** Performance in Predicting the Direction of Accessibility Difference of the best mouse-only vs. 3-species vs. 5-species models

| Species | Group | Metric | Mouse-Only | 3-Species | 5-Species | Number of Peaks |
|---|---|---|---|---|---|---|
| Cow | Val2 | Same Sign Count | 200 | 211 | 193 | 330 |
| Cow | Val2 | Same Sign (frac) | 0.606 | 0.639 | 0.585 | |
| Cow | Val2 | Pearson | 0.259 | 0.355 | 0.274 | |
| Cow | Val2 | Pearson P-Val | 3.80E-04 | 3.28E-09 | 8.33E-05 | |
| Cow | Val2 | Spearman | 0.263 | 0.388 | 0.279 | |
| Cow | Val2 | Spearman P-Val | 2.45E-04 | 2.54E-11 | 4.92E-05 | |
| Cow | Val2 | MSE | 0.399 | 0.656 | 0.584 | |
| Cow | Test2 | Same Sign Count | 358 | 364 | 365 | 618 |
| Cow | Test2 | Same Sign (frac) | 0.579 | 0.589 | 0.591 | |
| Cow | Test2 | Pearson | 0.201 | 0.34 | 0.309 | |
| Cow | Test2 | Pearson P-Val | 8.80E-05 | 3.32E-16 | 7.91E-13 | |
| Cow | Test2 | Spearman | 0.228 | 0.354 | 0.33 | |
| Cow | Test2 | Spearman P-Val | 1.85E-06 | 1.02E-17 | 8.14E-15 | |
| Cow | Test2 | MSE | 0.389 | 0.606 | 0.556 | |
| Macaque | Val2 | Same Sign Count | 308 | 311 | 313 | 520 |
| Macaque | Val2 | Same Sign (frac) | 0.592 | 0.598 | 0.602 | |
| Macaque | Val2 | Pearson | 0.269 | 0.318 | 0.321 | |
| Macaque | Val2 | Pearson P-Val | 9.60E-08 | 1.15E-11 | 1.36E-11 | |
| Macaque | Val2 | Spearman | 0.246 | 0.29 | 0.285 | |
| Macaque | Val2 | Spearman P-Val | 2.51E-06 | 1.53E-09 | 6.66E-09 | |
| Macaque | Val2 | MSE | 0.28 | 0.491 | 0.448 | |
| Macaque | Test2 | Same Sign Count | 466 | 464 | 467 | 798 |
| Macaque | Test2 | Same Sign (frac) | 0.584 | 0.581 | 0.585 | |
| Macaque | Test2 | Pearson | 0.212 | 0.241 | 0.245 | |
| Macaque | Test2 | Pearson P-Val | 2.74E-07 | 4.93E-10 | 4.21E-10 | |
| Macaque | Test2 | Spearman | 0.202 | 0.23 | 0.226 | |
| Macaque | Test2 | Spearman P-Val | 1.64E-06 | 4.41E-09 | 2.16E-08 | |
| Macaque | Test2 | MSE | 0.294 | 0.559 | 0.515 | |
| Pig | Val2 | Same Sign Count | 146 | 154 | 151 | 276 |
| Pig | Val2 | Same Sign (frac) | 0.529 | 0.558 | 0.547 | |
| Pig | Val2 | Pearson | 0.109 | 0.216 | 0.193 | |

| | | | | | | |
|---|---|---|---|---|---|---|
| Pig | Val2 | Pearson P-Val | 1.39E+01 | 3.06E-02 | 2.60E-01 | |
| Pig | Val2 | Spearman | 0.133 | 0.238 | 0.195 | |
| Pig | Val2 | Spearman P-Val | 5.45E+00 | 6.38E-03 | 2.26E-01 | |
| Pig | Val2 | MSE | 0.358 | 0.673 | 0.565 | |
| Pig | Test2 | Same Sign Count | 285 | 267 | 292 | 544 |
| Pig | Test2 | Same Sign (frac) | 0.524 | 0.491 | 0.537 | |
| Pig | Test2 | Pearson | 0.249 | 0.156 | 0.219 | |
| Pig | Test2 | Pearson P-Val | 7.55E-07 | 2.70E-02 | 4.79E-05 | |
| Pig | Test2 | Spearman | 0.195 | 0.124 | 0.18 | |
| Pig | Test2 | Spearman P-Val | 8.94E-04 | 3.90E-01 | 4.71E-03 | |
| Pig | Test2 | MSE | 0.343 | 0.755 | 0.616 | |
| Rat | Val2 | Same Sign Count | 483 | 470 | 470 | 796 |
| Rat | Val2 | Same Sign (frac) | 0.607 | 0.59 | 0.59 | |
| Rat | Val2 | Pearson | 0.159 | 0.289 | 0.27 | |
| Rat | Val2 | Pearson P-Val | 1.33E-03 | 9.23E-15 | 1.68E-12 | |
| Rat | Val2 | Spearman | 0.204 | 0.283 | 0.263 | |
| Rat | Val2 | Spearman P-Val | 1.34E-06 | 4.49E-14 | 8.47E-12 | |
| Rat | Val2 | MSE | 0.205 | 0.373 | 0.329 | |
| Rat | Test2 | Same Sign Count | 789 | 852 | 849 | 1440 |
| Rat | Test2 | Same Sign (frac) | 0.549 | 0.592 | 0.59 | |
| Rat | Test2 | Pearson | 0.177 | 0.293 | 0.285 | |
| Rat | Test2 | Pearson P-Val | 2.53E-09 | 5.84E-28 | 5.56E-26 | |
| Rat | Test2 | Spearman | 0.153 | 0.281 | 0.262 | |
| Rat | Test2 | Spearman P-Val | 1.17E-06 | 1.62E-25 | 1.10E-21 | |
| Rat | Test2 | MSE | 0.201 | 0.352 | 0.309 | |

P-values were adjusted using Bonferroni Correction. MSE: Mean squared error.
